# A framework for defining livestock ecotypes based on ecological modelling and exploring genomic environmental adaptation: the example of Ethiopian village chicken

**DOI:** 10.1101/2021.12.01.470795

**Authors:** Adriana Vallejo-Trujillo, Adebabay Kebede, Maria Lozano-Jaramillo, Tadelle Dessie, Jacqueline Smith, Olivier Hanotte, Almas Gheyas

## Abstract

In evolutionary ecology, an *ecotype* is a population that is genetically adapted to specific environmental conditions. Environmental and genetic characterisation of livestock ecotypes can play a crucial role in conservation and breeding improvement, particularly to achieve climate resilience. However, livestock ecotypes are often arbitrarily defined without a detailed characterisation of their agro-ecologies. In this study, we employ a novel integrated approach, combining Ecological Niche Modelling (ENM) with genomics, to delineate ecotypes based on environmental characterisation of population habitats and unravel the signatures of adaptive selection in the ecotype genomes. The method was applied on 25 Ethiopian village chicken populations representing diverse agro-climatic conditions. ENM identified six key environmental drivers of adaptation and delineated 12 ecotypes. Within-ecotype selection signature analyses (using *Hp* and *iHS* methods) identified 1,056 candidate sweep regions (SRs) associated with diverse biological processes. A few biological pathways were shared amongst most ecotypes and the involved genes showed functions important for scavenging chickens, e.g. neuronal development/processes, immune response, vision development, and learning. Genotype-environment association using Redundancy Analysis (RDA) allowed for correlating ∼33% of the SRs with major environmental drivers. Inspection of some strong candidate genes from selection signature analysis and RDA showed highly relevant functions in relation to the major environmental drivers of corresponding ecotypes. This integrated approach offers a powerful tool to gain insight into the complex processes of adaptive evolution including the genotype x environment (GxE) interactions.

## Introduction

Livestock genetic diversity is an essential prerequisite to achieve resilience to the challenges arising from climatic changes, and the ever-changing dynamics of production systems and consumer demands. Indigenous livestock populations surviving in diverse geographic areas exhibit unique genetic adaptations to their local environmental conditions. In evolutionary ecology, such locally adapted populations of a species are referred to as ‘ecotypes’. With their extant genetic and adaptive diversity, different ecotypes may hold genetic solutions for many present and future challenges facing the global livestock sector.

Delineating livestock ecotypes in practice, however, has been challenging. Arbitrary and loose criteria have often been applied for defining ecotypes, such as the geographic origin of populations, broad agro-climatic conditions of the populations or some major morphological features (Tadelle *et al*., 2003; Keambou *et al*., 2014; Sanarana *et al*., 2016). Such classifications are not only arbitrary but also are over-simplistic in that they neither take into consideration a detailed characterisation of the ecotype agro-ecologies nor do they attempt to decipher the key environmental drivers and their interactions in shaping ecotypes’ genomes. When a livestock species is introduced into a new environment, a wide range of bioclimatic conditions will interact with the standing genetic variations of the introduced population. Local agro-climatic conditions will exert different selection pressures in different environments and contribute to creating new ecotypes. This complex scenario is resonated in the ecotype defintion provided by Lowri (2012) who described the term as ‘a non-static adaptive variation over many traits across the natural landscape with no discernable boundaries’. This emphasises the need for adopting a ‘systems’ approach, such as environmental modelling, for defining ecotypes that would allow considering complex interaction of a large number of environmental predictors. Integrating such an approach with genomic characterisation will have major implications for the characterisation and conservation of livestock adaptive diversity and will help provide genetic solutions for developing ‘climate resilient’ livestock breeds.

Ecological Niche Modelling (ENM) – also called Species Distribution Modelling (SDM) - has been extensively applied for predicting distribution of wild animal species and crop plants. One of the most widely used ENM/SDM approaches is implemented in the program MaxEnt (Phillips *et al*., 2006). Using environmental data from known species-presence locations, MaxEnt predicts the potential geographical range of a species across a landscape and estimates the contribution of environmental variables in shaping the species’ habitats. As a consequence, the method has been a powerful tool in the conservation efforts of wild species (Thorn *et al*., 2009) and has found applications for other diverse purposes such as assessing the risk from invasive species (Jimenez-Valverde *et al*., 2011), epidemiological studies (Cardoso-Leite *et al*., 2014), and estimating the effect of climate change on future distribution of a species (Jeschke and Strayer, 2008). In livestock, however, the use of ENM is still in its infancy with only a few available examples (Pitt *et al*., 2016; Lozano-Jaramillo *et al*., 2018; Vajana *et al*., 2018; Gheyas *et al*., 2021; Kebede *et al*., 2021). Pitt *et al*. (2016) used MaxEnt for inferring the actual and historical distributions of early domestic fowl by modelling the environmental conditions of extant Red Jungle Fowl (RJF) with important conservation implications. Lozano-Jaramillo *et al*., (2018) used MaxEnt to predict the environmental suitability maps for successful introduction of two exotic chicken breeds across the Ethiopian landscape. Vajana *et al*., (2018) integrated landscape genomics and ENM to investigate local adaptation of indigenous Ugandan cattle to East Coast Fever. In a recent study (Gheyas *et al*., 2021), we applied MaxEnt for identifying the key environmental drivers in the agro-ecologies of Ethiopian indigenous chickens, followed by genomic analyses in relation to these variables to identify candidate adaptive genes. In another recent study, Kebede *et al*., (2021) integrated ENM with phenotypic distribution modelling to study phenotypic differentiation among Ethiopian indigenous chicken populations and used this characterisation for delineating potential ecotypes. The study, however, did not consider the genomic basis of adaptation.

Based on the insights from the above studies, here we present a novel application of ENM for delineating livestock ecotypes. Our approach considers ‘ecology’ as the main driving force for adaptive evolution and hence as the focal point for characterising ecotypes. The ENM-based ecotypes are then used as units of analysis to dissect the underlying genetics of adaptation. We exemplify the application of this approach using the same set of Ethiopian village chicken populations used in our previous study (Gheyas *et al*., 2021). Ethiopian indigenous chicken populations are an excellent model to demonstrate the utility of this novel approach for a number of reasons. In Ethiopia, indigenous chickens account for 78.85% of the total poultry population (CSA, 2021) and are predominantly reared in scavenging/semi-scavenging rural agro-ecologies, where the birds are directly exposed to various environmental pressures. A great deal of phenotypic diversity (Wilson, 2010) and genetic plasticity are observed in these indigenous chickens (Bettridge *et al*., 2018), even though their ancestral gene pool is not very diverse (Gheyas *et al*., 2021). Ethiopia’s agricultural landscape also varies widely due to its altitudinal topography and climatic variations, with one or two rainy seasons separated by dry seasons. As a result, the country displays diverse agro-climatic zones (n=18) (MoA, 1998). Indigenous chicken populations are found in most agro-ecologies where there is human settlement, indicating their adaptive diversity. The novel method introduced here offers an excellent platform for characterising these populations in relation to their environmental adaptation.

The major steps in our analysis include characterising the agro-ecologies of the investigated chicken populations using ENM, followed by clustering the populations based on their niche similarity to define distinct ecotypes. Within ecotype selection signature analyses are then performed to identify genomic loci at different stages of positive selection - ongoing and/or near fixation - using *iHS* (Integrated Haplotype Score) and *Hp* (Pooled heterozygosity) approaches. Variants overlapping the putative selective sweep regions are then analysed using Redundancy Analysis (RDA) approach to find their correlation with key environmental variables.

## Materials and Methods

### Study samples and environmental data

The data set included in this analysis comprised 245 chicken samples (Gheyas *et al*., 2021). Sampling was carried out as part of the African Chicken Genetic Gains project (https://africacgg.net/) to represent diverse agro-climatic conditions in Ethiopia. The samples originated from 25 villages or Kebeles across 12 districts and six national regions (Supplementary Table S1). For each population, 10 geographic coordinates, separated by 1.2 km^2^, were used to allow ENM-based characterisation of population habitats (see Gheyas *et al.,* 2021 for details).Thirty four environmental variables - 21 climatic, 1 elevation, 8 soil, and 4 vegetation and land cover conditions - were initially chosen for ENM as these were deemed important for chicken biology. Geo-referenced environmental data for these variables were extracted from public databases - viz. WorldClim (Fick and Hijmans 2017), SoilGrids (Hengl, *et al.,* 2014), Spatial Data Access Tool of NASA (ORNL_DAAC 2017), Harmonized World Soil Database (Fischer *et al.,* 2008) and Global Food Security-Support Analysis Data (GFSAD30 2017) - at a spatial resolution of 30 seconds (∼1 km^2^). Based on geodetic datum WGS84, the grids’ dimension and extension were corrected and homogenized for 1 km^2^ using ‘rgdal’, maptools’, rgeos’, and ‘raster’ packages in RStudio v1.1.419.

**Table 1.** Genome-wide pool heterozygosity (Hp) and integrated haplotype score (iHS) for the 12 proposed chicken ecotypes in Ethiopia.

| Ecotype | No. of Samples | Pool heterozygosity (*Hp*) statistics |  |  |  | Integrated haplotype score (*iHS*) statistics |  |  |  | Common overlapping candidate windows |
| --- | --- | --- | --- | --- | --- | --- | --- | --- | --- | --- |
| | | No. of windows analysed* | Mean ± SD | No. of candidate windows (unique ecotype specific windows) | No. of sweep regions<sup>§</sup> | No. of windows* | Mean ± SD | No. of candidate windows (unique ecotype specific windows) | No. sweep regions<sup>§</sup> | |
| E1 | 37 | 93031 | 0.30 ± 0.05 | 108 (14) | 37 | 92654 | 0.813 ± 0.36 | 191 (67) | 44 | 0 |
| E2 | 20 | 93224 | 0.31 ± 0.05 | 115 (22) | 35 | 92928 | 0.797 ± 0.40 | 243 (161) | 57 | 4 |
| E3 | 20 | 93191 | 0.32 ± 0.05 | 250 (59) | 64 | 92911 | 0.804 ± 0.38 | 132 (56) | 33 | 8 |
| E4 | 16 | 93042 | 0.31 ± 0.05 | 127 (9) | 30 | 92627 | 0.809 ± 0.39 | 98 (54) | 29 | 0 |
| E5 | 20 | 93258 | 0.31 ± 0.05 | 146 (30) | 39 | 92918 | 0.816 ± 0.40 | 420 (178) | 83 | 7 |
| E6 | 43 | 92687 | 0.30 ± 0.05 | 167 (38) | 49 | 92364 | 0.811 ± 0.33 | 58 (35) | 24 | 9 |
| E7 | 20 | 93200 | 0.29 ± 0.06 | 7 (0) | 2 | 92740 | 0.788 ± 0.39 | 205 (130) | 47 | 0 |
| E8 | 18 | 93175 | 0.30 ± 0.06 | 64 (8) | 13 | 92713 | 0.808 ± 0.41 | 264 (144) | 65 | 1 |
| E9 | 19 | 93277 | 0.32 ± 0.05 | 267 (70) | 63 | 92935 | 0.818 ± 0.41 | 441 (263) | 85 | 12 |
| E10 | 10 | 93313 | 0.33 ± 0.05 | 279 (65) | 77 | 93090 | 0.815 ± 0.40 | 81 (44) | 28 | 0 |
| E11 | 10 | 93261 | 0.31 ± 0.06 | 59 (20) | 18 | 92971 | 0.752 ± 0.50 | 549 (435) | 196 | 0 |
| E12 | 10 | 93266 | 0.32 ± 0.06 | 141 (38) | 37 | 93009 | 0.791 ± 0.43 | 159 (102) | 52 | 0 |

\* For both *Hp* and *iHS* analyses windows with SNP count $\geq 10$ were analysed but for *iHS*, filtration on SNP minor allele frequency ( $\geq 10\%$ ) reduced the number of windows.
<sup>§</sup> Candidate (sweep) regions were determined after merging overlapping or adjacent windows.

### Ecological Niche Modelling (ENM)

Prior to ENM, several complementary approaches were applied to evalutate the properties and relationship of the environmental variables and their relative contributions in explaining chicken agro-ecologies. This included a normality test, a Spearman’s rank correlation, and Principal Component Analysis (PCA). MaxEnt v3.4.1 (Phillips *et al*., 2006) was used for executing ENM. Model optimization was performed by removing highly correlated (Spearman *r* >0.6) and low contributing (<4%) environmental variables using ‘MaxentVariableSelection’ (Jueterbock *et al.,* 2016) and choosing the best combination of model parameters – Regularization Multiplier (RM) and Feature Classes (FCs) – using ENMeval (Muscarella, et al. 2014), considering all populations together (see Gheyas *et al*., (2021) for details).

Based on the optimization process, 6 environmental variables were shortlisted, namely, minimum temperature of coldest month (bio6 - minTemp), precipitation seasonality (bio15 - precSeasonality), precipitation of wettest quarter (bio16 - precWQ), precipitation of driest quarter (bio17 - precDQ), soil organic carbon (SoilOrgC), and land use (LandUse). The best combination of model parameters included Hinge (H), Quadratic (Q) and Product (P) FCs and RM = 3.5. A sub-sample of 25% was used as test data while the remaining 75% was used as training data for model execution. Habitat suitability maps were created in the logistic scale.

### Assessing niche similarity between populations and delineating ecotypes

Using the shortlisted variables and the optimal model parameters, habitat suitability maps were generated for individual populations. Similarity between suitability maps were assessed in ENMtools (Warren *et al*., 2010) using two methods: (i) pairwise Pearson correlation (*r)* and niche overlap statistic “*I*”. The *r* co-efficients capture correlation of corresponding grids between two suitability maps, where +1 indicates total positive linear correlation, 0 is no correlation, and -1 indicates complete negative correlation. Contrarily, the ‘*I’* values ranges from 0 indicating no overlap to 1 suggesting identical niche models between corresponding grids (Warren, 2008). The populations were grouped based on their habitat similarity by hierarchical clustering using the R package ‘cluster.’ Both values (*‘I’* and *‘r’*) were converted into “Euclidean” distance. Different hierarchical clustering approaches (minimum and maximum linkage, UPGMA, and Ward method) were evaluated to measure the cluster strength based on their agglomerative coefficients. Dendrograms and heatmaps for each similarity dataset were produced using the ‘gplots’ R package. Population clusters generated by these methods were considered as ecotypes when all population pairs within the cluster showed both the *I* and *r values* ≥ 0.6.

### Genome sequencing, SNP calling and genetic structure of the studied populations

Whole-genome sequencing of individual samples was performed on an Illumina HiSeqX platform with an average coverage of ∼45X . Sequence data processing, mapping, and variant calling have been detailed in Gheyas *et al.,* (2021).

Several analyses were performed to summarise the genetic diversity and population structure, as presented in Gheyas *et al.,* (2021). The previous results showed a mean nucleotide diversity between 0.28 and 0.34 per population, very low levels of population differentiation (based on pairwise Fst), a weak genetic sub-structure across populations (based on PCA analysis using SNP data) and the contribution of three different ancestral gene pools (based in Admixture analyses).

### Selective sweep analysis

Selection signature analyses were carried out on the ENM defined ecotypes by calculating pooled heterozygosity (*Hp)* following the method described by Rubin *et al*., (2010) and integrated haplotype score (*iHS)* using the Hapbin package (Maclean *et al.,* 2015). Both metrics were calculated in overlapping sliding windows of 20 kb with a step size of 10 kb.

For *iHS* calculation, phasing was performed (after removing missing genotypes) using Beagle v5.1 (Browning and Browining, 2007). Chromosome-specific mean recombination rates (cM/Mb) based on Groenen *et al*., (2009) or Eleferink *et al*., (2010) (for Chr16) were used for calculating genetic map positions of SNPs. Since no information was available for Chr30-33, a mean rate of 5.4 cM/Mb from across other micro-chromosomes (based on Groenen *et al*., 2009) was used. *iHS* analyses were first performed for individual SNPs by setting the minor allele frequency (maf) option as 10% and cut-off value for Extended Haplotype Homozygosity (*EHH)* equal to 0.1. Afterward, mean values were calculated within windows for standardised *iHS* (*iHS_std*) and the absolute value of *iHS_std*.

Empirical *P*-values were calculated for both standardised *Hp (ZHp)* and *iHS (iHS_std)* by ranking the windows in descending order according to their scores and then dividing the ranks by the total number of windows. The criteria for selecting outlier windows (i.e., potential sweeps) for *iHS* analysis included: windows with at least10 SNPs, *P*-value ≤ 0.01, window mean of |*iHS_std*| ≥ 2.0, and at least 90% of SNPs in a window with |*iHS_std*| value ≥ 2.0. For *Hp* analysis, the criteria were: windows with at least10 SNPs, *P*-value ≤ 0.01, and *ZHp* ≤ -4.0.

### Genotype-environmental association

Genotype-environment association analysis was performed with an LD-pruned set of SNPs overlapping the sweep regions (from *iHS* and *Hp analyses)* using Redundancy Analaysis (RDA) in Vegan v2.5-4 (Oksanene, 2015) following Forester (2019). LD pruning was performed with PLINK v1.9 (Purcell *et al.,* 2007), using the following parameters: --indep-pairwise, window-size = 10 kb, step-size = 10 SNPs, and r^2^ = 0.5. Genotype was fitted as a response variable and environmental data (6 variables) as the explanatory variables with conditioning on latitude and longitude. Ancestry co-efficients from Admixture analysis were included as covariates to correct for any population structure effect.

For each canonical RDA axis, ANOVA analysis was performed. The significance of the partial model and each axis was calculated with 999 and 99 permutations, respectively.

### Functional annotation

Candidate regions/variants from selection signature and RDA analyses were investigated for their overlap with chicken genes from Ensembl (release 98). SNPs were annotated using Variant Effect Predictor (VEP) (McLaren *et al.,* 2016). Candidate genes were checked for their overlap with chicken-QTLs from AnimalQTLdb (release 45) (Hu *et al.,* 2019). Only QTLs with *P*-value <0.05 and size <1Mb were considered. Molecular functions, biological processes, and metabolic pathways associated with the candidate genes were analysed using the Panther classification system v.14.0 (Huaiyu *et al.,* 2019).

## Results

An overview of our framework for ENM-based delineation of livestock ecotypes and their use for dissecting genomic-environmental adaptation is presented in Figure 1 with summary results from its application on Ethiopian chickens.

**Figure 1.**
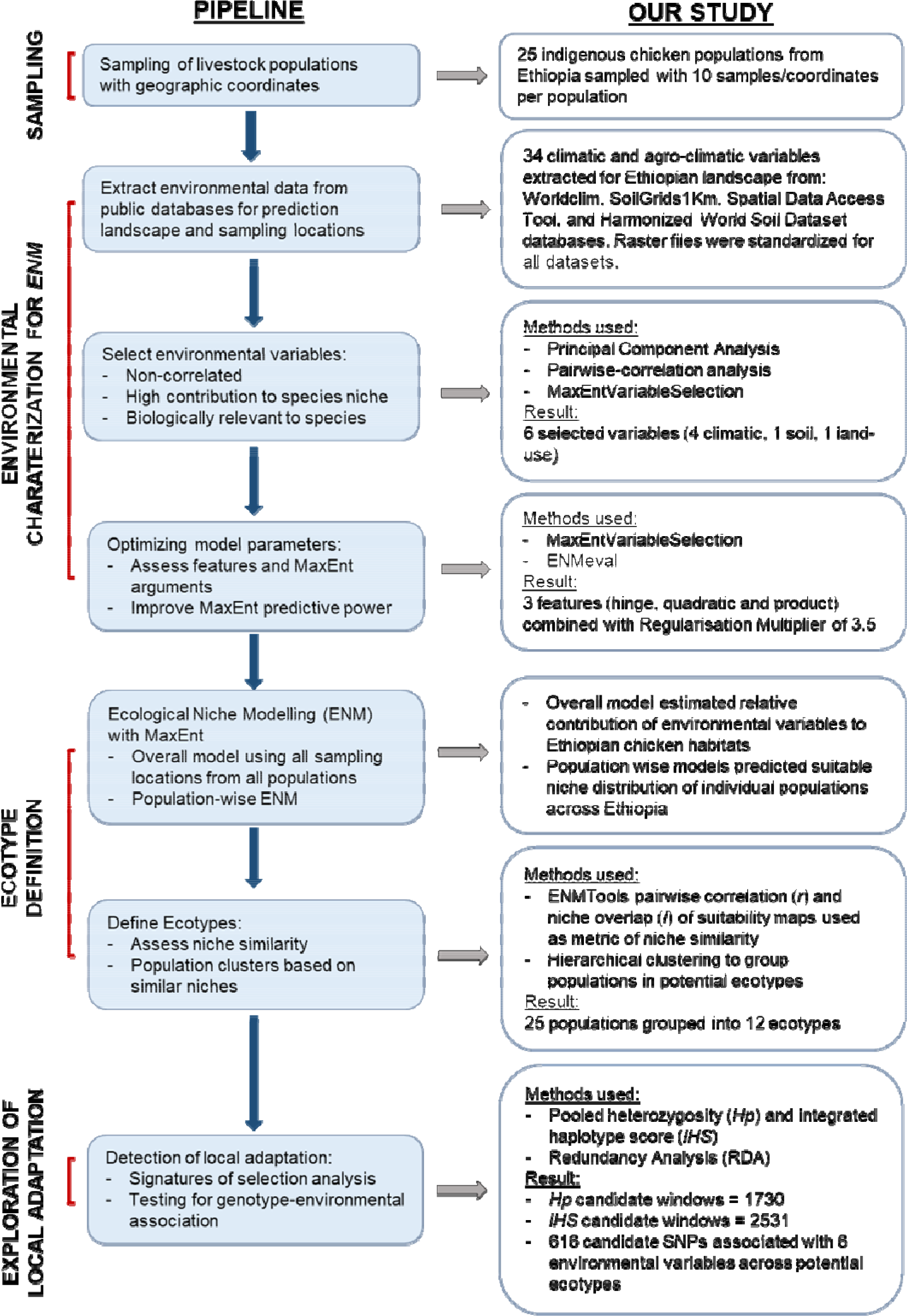
A framework for defining livestock ecotypes based on ENM approach and exploring genomic adaptation to the environment.

### Established agro-ecological zones (AEZ) are not sufficient for classifying populations into ecotypes

Different classifications of Ethiopian AEZs are available. Since altitudinal topography constitutes a major feature of the Ethiopian landscape, traditional classifications were driven by this and 4-6 different AEZs were defined, e.g., Berha (lowlands, <500 m.a.s.l.), Kolla (lowlands, 500 - 1,500 m.a.s.l.), Weyna Dega (midlands, 1,500 -2,300 m.a.s.l.), Dega (highlands, 2,300 -3,200 m.a.s.l.), Wurch (highlands, 3,200 – 3,700 m.a.s.l.), and High Wurch (highlands, >3,700 m.a.s.l.) (Hurni 1998). Many other classifications have combined rainfall and crop pattern information with these traditional elevation-based zonation for the purpose of crop management, but their denominators were often relative and lacked precision (Hurni 1998). The FAO/IIASA-defined ‘Global AEZ 16-class’ classification (HarvestChoice/IFPRI 2009) has also been applied on Ethiopia and identifies 8 different zones (Figure 2A). While this classification considers temperature, elevation, and rainfall, still it suffers from lack of precision. For example, it provides only two broad classifications based on temperature - either cool or warm.

**Figure 2.**
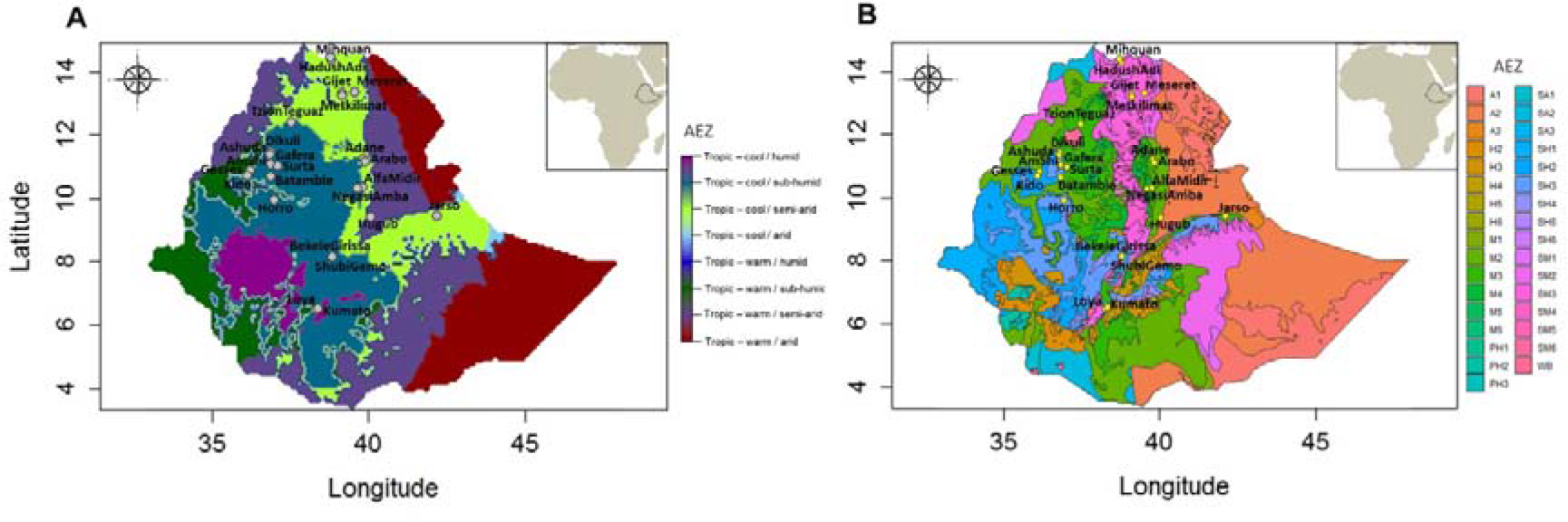
Ethiopian agro-ecological zones and sampling locations. **(A) Population locations against Global 16-Class AEZ classification (courtesy of Gheyas et al., 2021), and (B) population locations against the elaborate agro-ecological zonation by MoARD**. A1=Hot arid lowland plains; A2=Warm arid lowland plains; A3= Tepid arid mid highlands; SA1= Hot semi-arid lowlands; SA2: Warm semi-arid lowlands; SA3: Tepid semi-arid mid highlands; SM1= Hot sub-moist lowlands; SM2 = Warm sub-moist lowlands; SM3=Tepid sub-moist mid highlands; SM4=Cool sub-moist mid highlands; SM5=Cold sub-moist mid highlands; SM6=Very cold sub-moist mid highlands; M1=Hot moist lowlands; M2=Warm moist lowlands; M3=Tepid moist mid highlands; M4=Cool moist mid highlands; M5=Cold moist sub-afro-alpine to afro-alpine; M6=Very cold moist sub-afro- alpine to afro-alpine; SH1=Hot sub-humid lowlands; SH2=Warm sub-humid lowlands; SH3=Tepid sub-humid mid highlands; SH4=Cool sub-humid mid highlands; SH5=Cold sub- humid sub-afro-alpine to afro-alpine; SH6=Very cold sub-humid sub-afro alpine to afro- alpine; H2=Warm humid lowlands; H3=Tepid humid mid highlands; H4=Cool humid mid highlands; H5=Cold humid sub-afro-alpine to afro- alpine; H6=Very cold humid sub-afro- alpine; PH1=Hot per-humid lowlands; PH2=Warm per-humid lowlands; PH3=Tepid per- humid mid highland; WB= waterbody.

The most elaborate zoning is from Ethiopia’s Ministry of Agriculture and Rural Development (MoARD, 2005) with the identification of a staggering 32 different AEZs (https://www.yieldgap.org/ethiopia) (Figure 2B), taking into account climate, physiography, soil properties, vegetation, and crop patterns. Although this provides a high-resolution classification, the system is developed mainly for agricultural crop management and therefore may not be directly relevant for livestock species.

The MaxEnt based ecological modelling (ENM) for characterising livestock agro-ecologies offers a number of advantages over these established classifications. ENM provides the opportunity to incorporate species-relevant environmental variables and allows examination of a large number of variables. In our study, we initially examined 34 different variables (Figure 3A). Elevation and climatic parameters representing temperature, rainfall patterns, and atmospheric humidity were included, as these may challenge the physiological tolerance of chickens. Soil property variables were included as these could affect food availability for scavenging birds. Vegetation and land use variables were incorporated due to their potential effects on the type of food chickens have access to, as well as their role in providing shelter from predators and inclement weather. Figure 3A shows the relative contribution of these variables in explaining Ethiopian chicken agro-ecologies and marks the most important uncorrelated variables selected by MaxEntVariableSelection for final modelling. ENM’s ability to examine individual variable contribution is extremely important for gaining insight into which variables may play key roles in driving adaptation. Shortlisting of the variables on the other hand is important to avoid ‘overfitting’ of the models and to remove collinearity from the model predictors that can otherwise increase model uncertainty and reduce efficiency (De Marco and Nóbrega, 2018).

**Figure 3:**
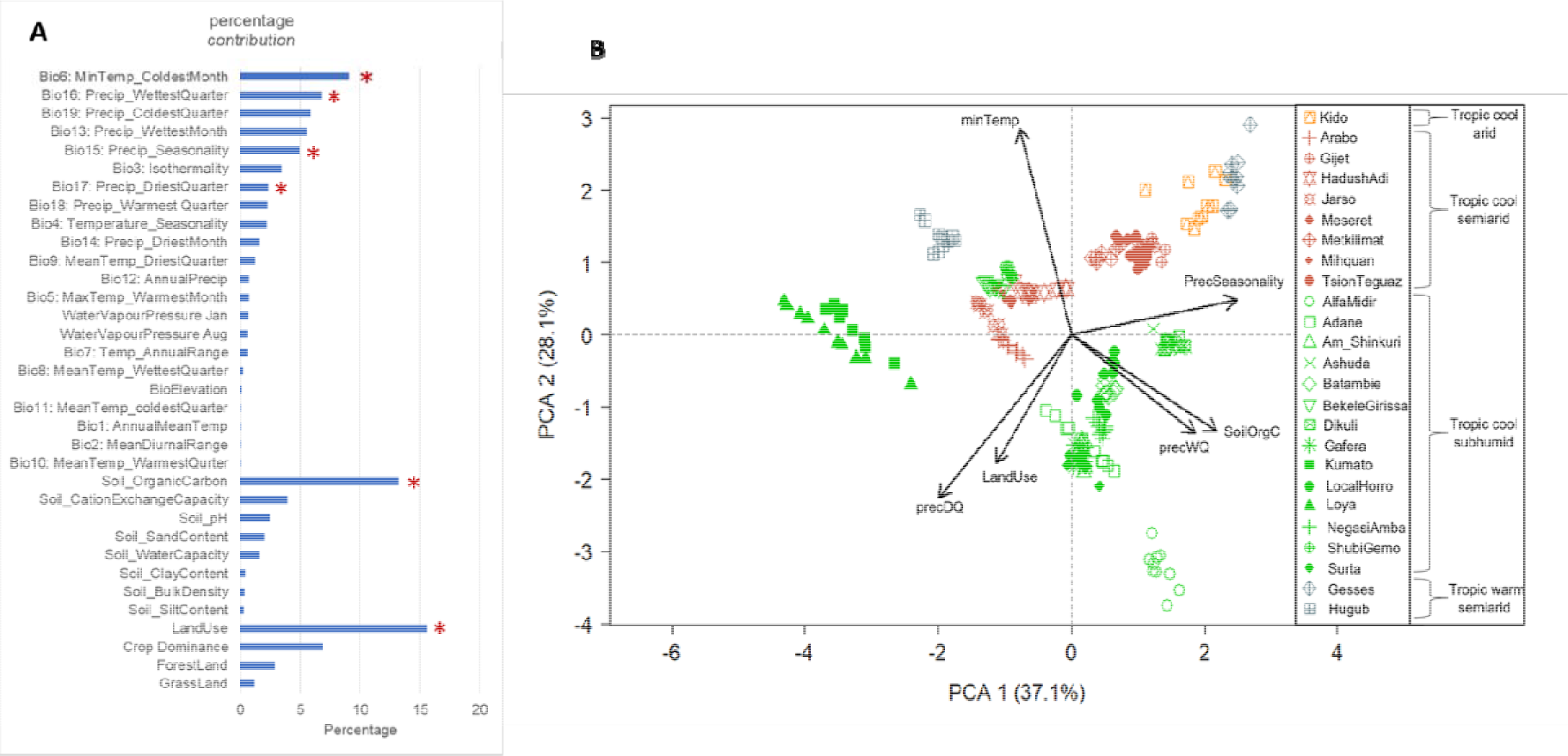
Environmental *variables* used in ENM for characterising Ethiopian chicken agro-ecologies. (A) Relative contribution of the 34 agro-climatic variables examined in Ethiopian chicken agro-ecologies. Asterisks mark the variables that were shortlisted for final modelling. (B) PCA plot of populations based on 6 shortlisted variables. The colour coding shows the population classification based on Global AEZ-16. Arrow direction shows the contributing variables in spreading the samples and arrow length indicates relative contribution.

Figure 3B compares the partitioning of the populations based on the six key environmental factors selected by ENM to FAO/IIASA AEZ classification. Some populations clustered together in the PCA plot are from different FAO/IIASA AEZs (e.g., Kido from the cool-arid zone and Gesses from the warm-semiarid zones or Bekele Girissa and Shubi Gemo from the cool-subhumid and Jarso, Hadush Adi and Mihquan from the cool-semiarid zones). Others from the same AEZ are far apart in the PCA plot. For example, 14 of the 25 populations belonging to the broad cool-subhumid AEZ are split into 3-4 major clusters (e.g. Loya, Alfa Midir, and Ashuda, with different environmental parameters playing key roles in their habitat stuctures). This signifies the importance of ENM in characterising the agro-ecologies and identifying the key environmental drivers in population habitats.

### ENM characterises Ethiopian village chicken agro-ecologies

As described in the Methods section, MaxEnt models were executed first on all populations combined and then on individual populations using the six selected variables and the best parameter settings. The predictive power of all the models, as assessed by AUC (Area Under ROC Curve) values, were high (>0.99) for both training and test data. Similarly, the AUC values for the individual environmental variables for each population were generally higher than 0.6, with several exceptions, like bio6 (minTemp) and bio15 (precSeasonality), which in some populations had values less than 0.6 but greater than 0.5, which still represent moderate potential in predicting suitable conditions (Supplementary Table S2).

**Table 2.** Selected RDA SNPs outliers overlapping protein-coding genes.

| Outlier SNP | Gene | Ecotype* | Gene function / association | Reference |
| --- | --- | --- | --- | --- |
| **MinTemp** |  |  |  |  |
| 4:88536330 | *PANK2* | E11 | Neuronal and vascular development; aerobic respiration | Zizioli *et al.*, (2016); Uniprot |
| 2:78280162 | *MARCHF6* | E10 | Involved in protein ubiquitination pathway; metabolic integrator in cholesterol synthesis. | Scott *et al.*, (2021) |
| **PrecSeasonality** |  |  |  |  |
| 1:71668364 | *LRP6* | E2 | Involved in Wnt Signalling pathway. | Chao *et al.*, (2019) |
| 10:9402417 | *MAPK4* | E11 | Found associated with early body temperature and *Salmonella pullorum* resistance in chickens. | Li *et al.*, (2020) |
| 4:84643310 | *CTBP1* | E11 | Co-repressor targeting diverse transcription regulators | Uniprot |
| 5:19554343 | *LDLRAD3* | E12 | Association with heat stress and Newcastle Disease Viral (NDV) infection in chicken | Jastrebski *et al.*, (2017); Wang *et al.*, (2020) |
| **precWQ** |  |  |  |  |
| 2:117749087 | *RDH10* | E2 | Retinoid (Vitamin A) metabolic process | Uniprot; D'ambrosio *et al.*, (2011); |
| 10:13599109 | *DET1* | E11 | Protein ubiquitination and degradation; associated with salt and osmotic stress in plant. | Uniprot; Fernando *et al.*, (2018) |
| 21:4353990 | *FBXO42* | E8 | Protein ubiquitination and degradation | Uniprot |
| 4:3627776 | *HS6ST2* | E10 | Associated with growth and carcass traits in chicken. | Wang *et al.*, (2018) |
| 4:82439583 | *TNIP2* | E11 | Involved in stress-activated MAPK cascade and immune and inflammatory response. Detected in Ugandan chicken ecotypes in association with environmental stress. | Uniprot; Fleming *et al.*, (2016) |
| **precDQ** |  |  |  |  |
| 1:197483358 | *DCHS1* | E7 | Neurogenesis, cardiac valve formation | Uniprot |
| 20:1187011 | *ERGIC3* | E4, E7 | Cytoplasmic transport protein; plays role in ER stress-induced cell death and in cell growth. | Uniprot; OMIM |
| 8:24798255 | *GPX7* | E7 | Cellular response to oxidative stress; found associated with heat stress in chickens and livestock and draught stress in plants. | Del Vesco *et al.*, (2015); Berihulay *et al.*, (2019); Cruz de Carvalho (2008) |
| 19:1816989 | *AUTS2* | E8 | Neurological development. | Uniprot |
| **SoilOrgC** |  |  |  |  |
| 1:23065714,<br>1:23140159,<br>1:23141490,<br>1:23164664 | *PTPRZ1* | E7 | Central nervous system development; associated with learning or memory | Uniport |
| 5:19564424 | *LDLRAD3* | E12 | Associated with lifespan extension in mice | Tyshkovskiy *et al.*, (2019) |
| 19:1819556,<br>19:1830033 | *AUTS2* | E8 | Neurological development. | Uniprot |
| 10:7190022 | *MYO1E* | E11 | Intracellular movements and membrane trafficking. | Uniport |
| 19:1819556 | *LMF1* | E8 | Triglyceride metabolic process | Uniprot |
| 3:16891572 | *GPCPD1* | E12 | Skeletal muscle tissue development; Differential expressed in modern pedigree broiler line compared to unselected broiler line. | Kong *et al.*, (2017) |
| 3:38560061 | *TARBP1* | E6 | Transcription regulation; tRNA methylation | Uniprot |
| 5:35557653 | *NPAS3* | E7 | Neurodevelopment; regulation of glucose metabolism | Sha *et al.* (2012) |
| 5:39594296 | *ADCK1* | E8 | Involved in mitochondrial functions; lipid homeostasis | Uniprot |
| 6:28550216 | *NHLRC2* | E12 | Required for normal embryonic development | Uniprot |
| 7:30965068,<br>7:30967748 | *THSD7B* | E12 | Candidate gene for albumin quality in chicken | Qu *et al.*, (2019) |
| 8:6198062 | *STX10* | E12 | SNARE protein involved in intracellular protein transport. | Uniprot |
| **LandUse** |  |  |  |  |
| 17:7056075 | *GFI1B* | E11 | Transcription factor with important role in regulation of hemopoiesis. | Uniprot |
| 17:7239853 | *SLC2A6* | E11 | Regulation of glycolytic process; glucose transporter activity | Uniport |
| 2:31712297 | *OSBPL3* | E4 | Regulates hepatic fat metabolism, involved in insulin signalling pathways | Yan *et al.*, (2007) |

\* The ecotypes from which the SNPs originally overlapped with sweep regions

The combined analysis of populations allowed assessment of the effect of individual variables on the predictive power of the model (Figure 4). The jackknife result of test and training gains is an important metric for assessing variable contribution and model performance (Figure 4A and 4B). Jackknife results indicated that, in our model, bio6 (minTemp), bio16 (precWQ), LandUse and soilOrgC were the variables that had the most useful information when used in isolation of other predictors. On the contrary, bio15 (precSeasonality) decreased the gain most when excluded from the analysis, indicating that this variable contained the most information that was not present in any other predictors.

**Figure 4:**
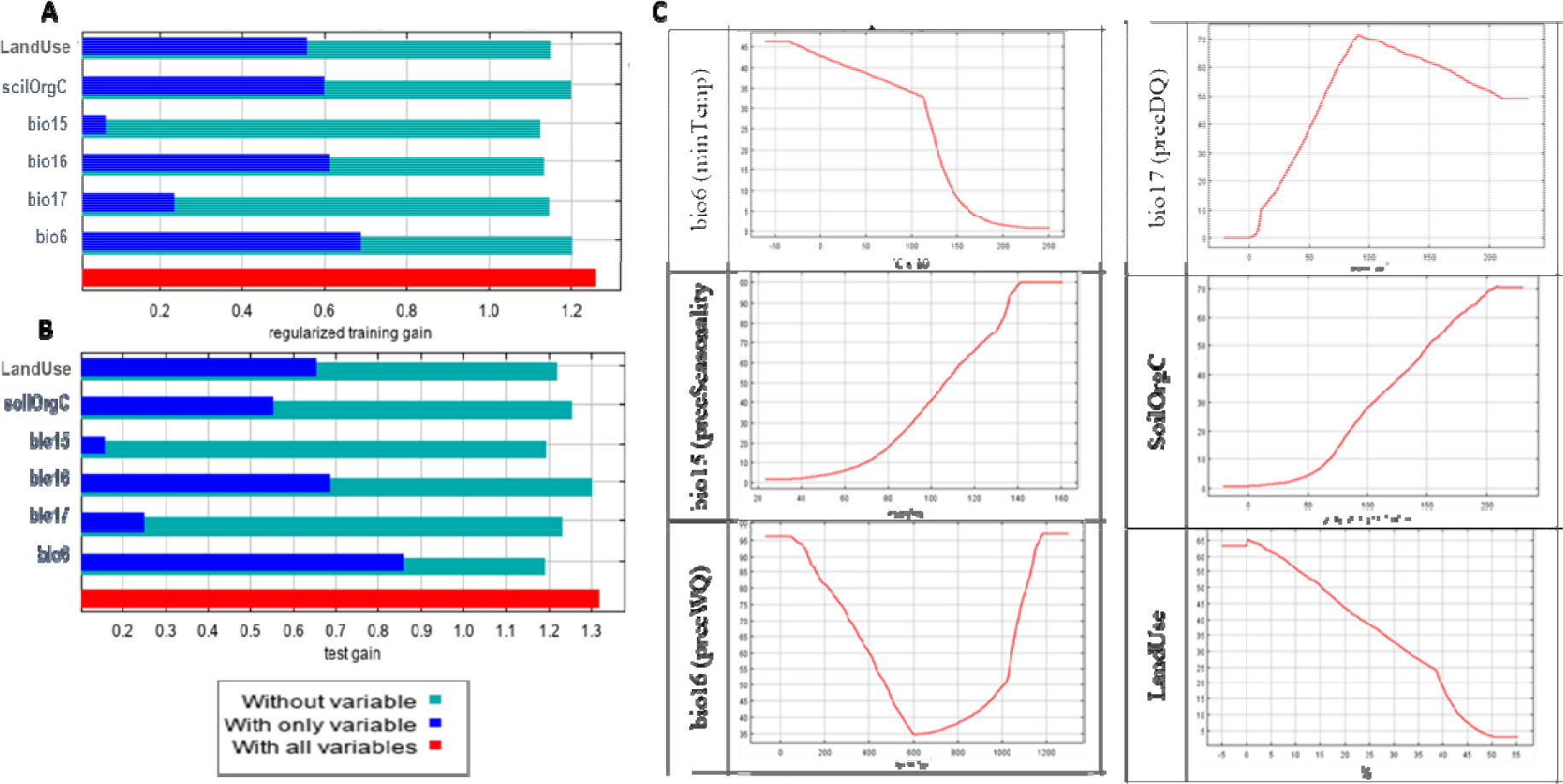
Predictive powers of individual environmental variables in ENM. Results from the combined model based on all 25 populations using 6 selected environmental predictors showing (A) Jacknife of training gains for models with or without a variable, (B), Jacknife of test gain for models with or without a variable, and (C) marginal response curves for individual variables. The X-axis in the response plots shows the environmental values and Y-axis shows the logistic output for probability of presence.

We also inspected the marginal response curves (Figure 4C) from the overall model to examine how the predicted suitability varies as each environmental variable changes while keeping all the other variables at their average sample value. The curves for minTemp and LandUse show that the probability of suitability decreases with rise in these parameters . On the other hand, when precSeasonality and soilOrgC increase, the likelihood of chicken presence increases. For precWQ, the response curve drops sharply with increase in the variable’s value until 600 mm/m^2^ but then rises sharply again, indicating its potential interaction with other environmental variables. In contrast, the likelihood of chicken presence increases when precipitation in precDQ rises to 90 mm/m^2^ and drops slowly until 200 mm/m^2^.

In the combined population model, the contributions of the six variables ranged between 10% for LandUse and 24% for SoilOrgC. The remaining 66% of contributions came from climatic variables: minTemp (21%), precWQ and precSeaonality (both 16%), and precDQ (13%). The relative contributions of the variables in different populations, however, varied widely as shown in Supplementary Figure S1 and not all variables contributed to the characterisation of every population agro-ecology. Supplementary Figure S2 shows the environmental suitability maps for individual populations across the Ethiopian landscape.

### Delineating ecotypes based on niche similarity

Pearsons correlation (*r*) and Niche Overlap (*I*) statistics were generated by pairwise comparison of the suitability maps for clustering the populations with similar niche for defining ecotypes (Supplementary Tables S3a and S3b). These two metrics provide complementary information. The Niche Overlap metric is a measure of similarity of ocurrences between two populations. In contrast, correlation statistic compares the underlying models (Warren and Seifert, 2010). The correlation metric is generated by comparing all corresponding grid cells between two maps. On the other hand, the Niche Overlap statistic only checks for the proportion of overlap of grid cells that were predicted as suitable (i.e. with logistic suitability score = 1) between two maps. In our study, the results of these two metrics were significantly correlated (Spearman correlation coefficient = 0.6; *P* < 0.0001).

Among the different hierarchical clustering approaches evaluated, the Ward method was selected as it had the largest agglomerative coefficient (0.746). Clustering of the populations based on the above two similarity metrics showed different overall topography (Figures 5A and 5B). For delineating ecotypes, we considered the clustering based on one or both approaches as a guide but the definitive criterion was that all pairwise comparisons within an ecotype showed niche overlap *(I)* ≥ 0.6 and correlation co-efficient *(r)* ≥ 0.6.

**Figure 5:**
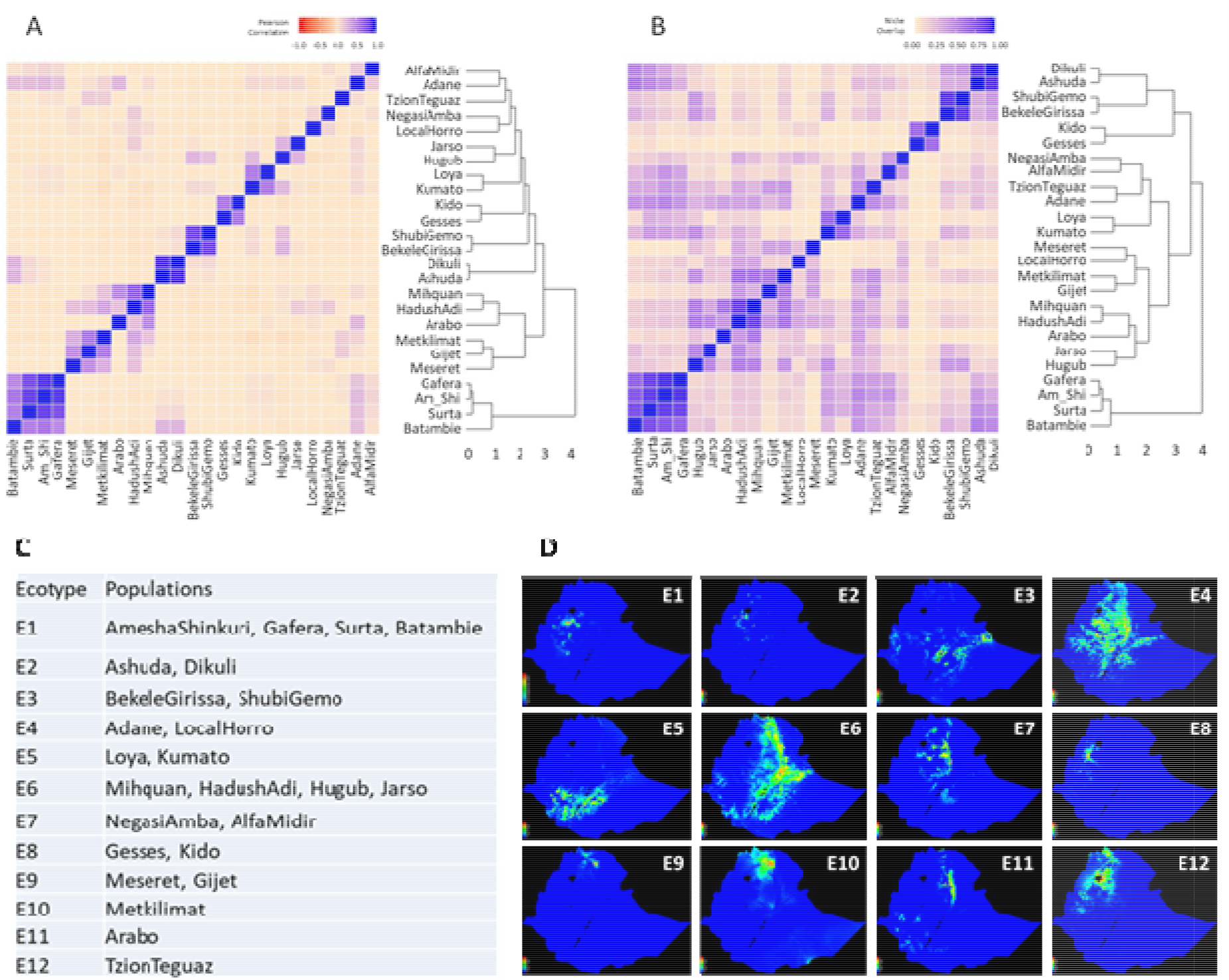
Delineation of ecotypes based on niche similarity among 25 Ethiopian chicken populations examined. (A) Dendrogram and heatmap of pairwise Pearson correlation *(I)* among 25 populations; (B) dendrogram and heatmap of pairwise niche overlap statistic *(r)* among 25 populations; (C) 12 delineated ecotypes and their population composition; (D) Suitability maps of 12 delineated ecotypes.

Both approaches grouped identically 16 out of the 25 populations. Consistent clusters were:

(1) Amesha Shinkuri, Gafera, Surta, and Batambie; (2) Ashuda and Dikuli; (3) Bekele Girissa and Shubi Gemo; (4) Gesses and Kido; (5) Kumato and Loya; (6) Hugub and Jarso, (7) Hadush Adi and Mihquan. The first five clusters were used to define individual ecotypes whereas the last two clusters were merged into a single ecotype, as both niche similarity matrics fulfilled the criterion of ≥ 0.6 for all population pairs after merging.

For the remaining 9 populations, an examination of (*r*) and (*I*) values was performed to resolve their ecotype membership. Only three populations – Arabo, Metkilmat, and TzionTeguaz - could not be grouped with any other populations and hence were considered as separate ecotypes. Thereby, 12 ecotypes were identified for the 25 populations studied (Figures 5C and 5D), reflecting the large heterogeneity of the Ethiopian agro-ecological landscape. Notably, even though many populations belong to the same broad AEZ, multiple ecotypes were delineated from them based on ENM, e.g., the cool/sub-humid zone overlaps ecotypes E1-5 and E7 (Supplementary Table S1). Contrarily, populations in E6 originate from two broad AEZs - cool/semi-arid and warm/semi-arid. Although in most cases, geographically close populations were clustered into ecotypes (E1-E3, E5, E7-E9), that did not hold true in all cases. For instance, the E4 populations – Adane and Horro – are far apart geographically (∼300 km). Similarly in E6, Mihquan and Hadush Adi populations are geographically close (∼40 km) but are far from Hugub and Jarso (∼570 - 660 km). In contrast, Adane and Arabo are geographically close (∼10 km) but were placed in separate ecotypes. These results demonstrate the large heterogeneity in environmental conditions in Ethiopia even within a short geographic distance, and thereby shows the importance of ENM-based characterisation of agro-ecologies. The relative contributions of the 6 environmental variables in the 12 ecotypes are presented in Supplementary Figure 3.

### Genomic analyses identify candidate selective sweep regions within ecotypes

About 14M autosomal SNPs detected in Gheyas *et al*. (2021) were used for selection signature analyses in the present study. Genetic variant data from all populations constituting each ENM-defined ecotype were combined, resulting in 11M to 13M SNPs per ecotype. Selection signature analyses were then performed within each ecotype using *Hp* and *iHS* approaches.

The genome-wide mean values of *Hp* were similar across all the ecotypes, ranging from 0.29 ± 0.06 for E7 and 0.33 ± 0.05 for E10 (Table 1). However, the number of *Hp* candidate windows varied widely from only 7 for E7 to 279 for E10. Adjacent or overlapping candidate windows were merged to define the sweep regions (SRs). This resulted in 2 to 77 SRs per ecotype and a total of 365 regions from all ecotypes combined (Table 1, Supplementary Table S4).

The genome-wide mean values of *iHS* were also similar among the ecotypes, ranging from 0.75 ± 0.5 (E11) to 0.81 ± 0.4 (E9) but again the number of candidate sweep windows showed large variation, ranging from 58 (E6) to 549 (E11) (Table 1). Interestingly, although E10 produced the largest number of candidate signals with *Hp* analysis, this ecotype produced one of the lowest number of candidate windows (only 81) with *iHS* analysis. Merging the overlapping candidate *iHS* windows resulted in 24-196 SRs in different Ecotypes (Table 1). A complete list of unique SRs (n=1,056) from both methods can be found in Supplementary Table S4 along with overlapping genes. The SRs vary in size from 20 Kb to 360 Kb and together cover roughly 4% of the chicken genome.

### Shared sweeps between methods and among ecotypes

We found a weak negative correlation between *Hp* and *iHS* (-0.043 ± 0.06, *P*-value = 0.00), and *ZHp* and *iHS_std* (-0.18 ± 0.06, *P*-value = 0.00) values in sweep regions (Supplementary Figures S4a and S4b). This shows a complementarity of the methods which is expected as *Hp* identifies regions that are fixed or near fixation (Rubin *et al.,* 2010), whereas *iHS* detects ongoing selection (Voight *et al.,* 2006). Despite this, we found a few SRs (n = 17) which were commonly detected by both methods either within the same ecotype (n = 8) or from different ecotypes (Supplementary Table S5, Supplementary Figure S5). Regions commonly detected by both methods within the same ecotype, would indicate it is close to reaching fixation but not yet fixed. In contrast, the same SR detected by *Hp* in one ecotype and *iHS* in another, may pinpoint different levels of selection in different ecotypes.

The percentage of shared sweep windows among ecotypes varied from 1% to 13% (Supplementary Figure S6). E11 generally showed the lowest level of shared sweeps (1% - 2%) with any other ecotypes whereas E3 generally shared the largest proportion with many other ecotypes (e.g., it shared 10 to 13% of its sweep windows with five other ecotypes).

We also investigated the shared sweep region genes among ecotypes . About 22% (n =1. 125) of the genes overlapping *Hp-*based SRs were shared by at least two ecotypes (Supplementary Table S6a). The *TSHR* (Thyroid Stimulating Hormone Receptor) gene was detected ubiquitously in all ecotypes. This gene has previously been detected as a selection signal (Rubin *et al*., 2007; Gheyas *et al.,* 2015) and therefore represents an old sweep, possibly associated with chicken domestication (Rubin *et al*., 2007; Karlsson *et al.,* 2016). Several other *Hp* candidate genes were detected in the majority of the ecotypes, e.g., *ENSGALG00000047413* (11 ecotypes) and *ENSGALG00000052351* (10 ecotypes) – both LncRNAs with possible *cis*-acting regulation on nearby genes (e.g., *TSHR* and *FUT8*, respectively), *TSNARE1* (9 ecotypes) and *SCN2A* (8 ecotypes) – both neurobehavioral genes, and *CACNA2D3* (7 ecotypes) with voltage-gated calcium channel activity regulating ion transmembrane transport. In comparison to the *Hp* analysis, only 10% (122 of 1,180) of the *iHS*-genes were shared by 2 to 5 ecotypes at most (Supplementary Table S6b).

### Candidate sweeps are related to diverse biological functions and phenotypes

Genes overlapping the SRs from different ecotypes were checked for their functional classification according to Panther Pathways (Supplementary Figure S7). Although only 1,253 genes (∼5% of all chicken genes) intersected SRs from different ecotypes, they showed a hit for 55% (98 of 177) of the Panther pathways, indicating their involvement in a vast array of physiological processes. About 77% (75 of 98) of these pathways are represented by SR genes from multiple ecotypes but the number of genes contributed by the ecotypes varied. A few pathways are represented by genes from all or most ecotypes. For instance, *Alzheimer disease-amyloid secretase pathway* (*P00003*), and *Heterotrimeric G-protein signalling pathways* (*P00026* and *P00027*) had hits from all 12 ecotypes with representation by 1 to 6 genes per ecotype. Similarly, *Integrin signalling pathway (P00034) and PDGF signalling pathway (P00047)* have hits from 10 different ecotypes. Investigation of the individual genes from these ubiquitous pathways shows involvement in a myriad of biological processes, but prominently in processes which are expected to be necessary for the survival of scavenging chickens, irrespective of ecotypes; for instance, roles in nervous system development, neurological processes and/or cognitive functions (*APBA2, MAPK6, MAPK13, MAPK14, PCSK2, CACNA1C, BACE2, PKN2, GPSM1, MTNRIB, DRD3, HRH1, GRM5, GRB2, CRKL, MAP2K2, MICALL1, NTN4, RND2, FOS, MTOR*, *PRKAR2B, DLC1*); eye development, visual perception and visual learning (*MTNRIB, GRK7, DRD3, HRH1, COL5A1*) – important for foraging for food; immune response, inflammatory response, and wound healing (*MAPK14, RASGRP1, GRB2, HRH1, CREB3L3, CHUK, CRKL, FOS, MTOR*); and apoptotic process (*MAPK14, PKN2, DRD3, ACTN1, VAV2, DLC1*) – an important stress response (Uniprot; Genecards; Kawamata *et al*., 1998; Chen *et al.,* 2017; Yan, 2017; Huentelman *et al*., 2019; Asih *et al*., 2020). Other notable involvements include regulation of cardiovascular and renal development and functions (*CACNA1C, VAV2, MAPK14, DLC1, PCSK2, COL5A1, CRKL, MTOR, PRKAR2B, DRD3, NTN4, RND2, ADCY2*) (Uniprot).

About 12% (n=128) of the SRs overlapped with known chickens QTLs (Figure 6; Supplementary Table S7), indicating potential association with different phenotypic traits, e.g., reproductive efficiency (n=28 hits), egg quality (n=21), growth (n=20), feather pecking behaviour (n=14), feed conversion efficiency or FCR (n=12), immunity and health (n=11), body fat (n=10), body temperature (n=4), feed intake (n=4), and feather pigmentation (n=4).

**Figure 6:**
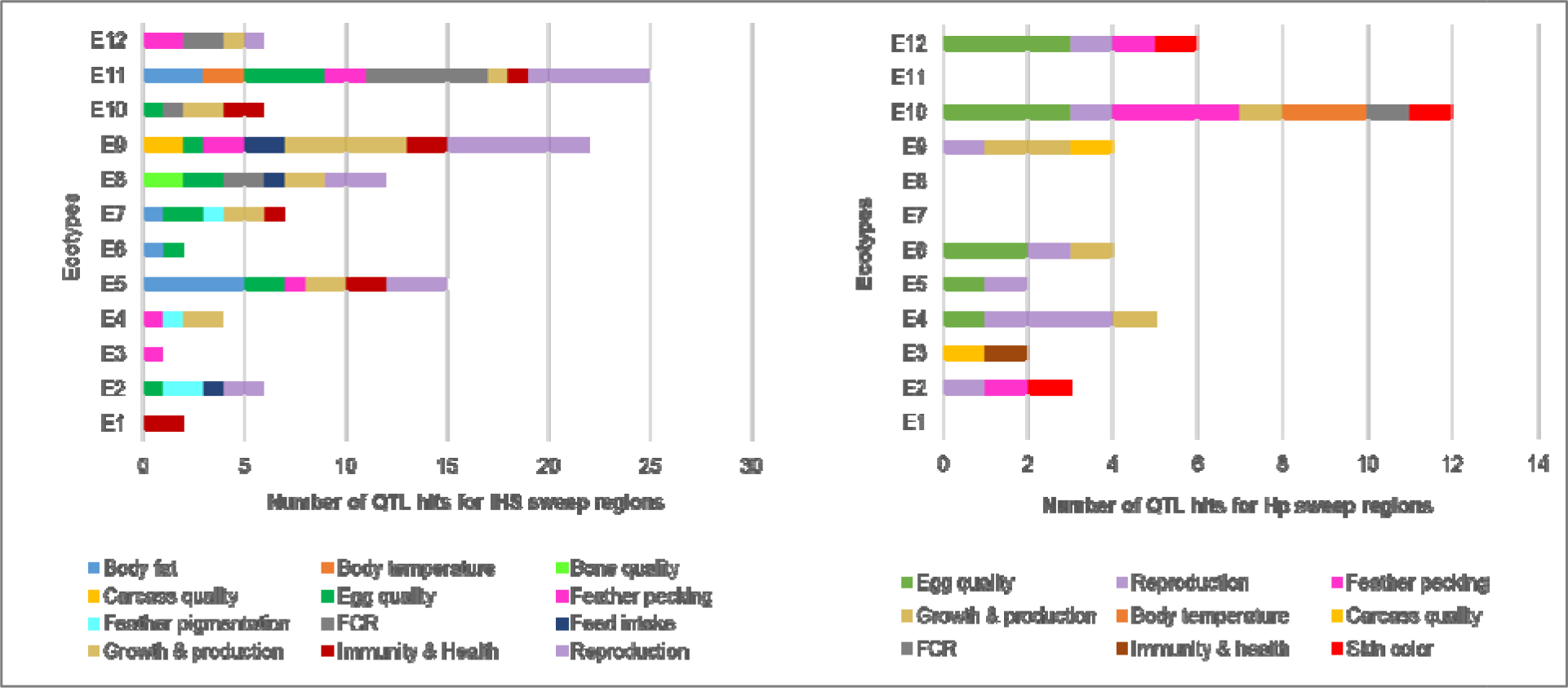
Overlap of the sweep regions from Hp and iHS analyses with chicken QTLs.

Finally we closely inspected the strongest signalling SR from each ecotype to see if a functional relevance can be drawn between the overlapping genes and the ecotype’s environment (Supplementary Tables S8a and S8b). Strikingly, in most cases a strong relevance is observed through the involvement of the genes in various stress responses, neuronal processes, immune responses and transcriptional regulations. The environmental association of the gene functions is more prominent for ecotypes where only a few environmental variables act as major driving forces. E7 perhaps offers the most striking example where the minTemp contributes ∼99% in characterising the ecotype’s agro-ecology. This ecotype has the lowest temperature environment (as low as near 0°C) due to its very high-altitude location (>3,000 m.a.s.l.). The strongest *iHS*-sweep from this ecotype overlaps several genes: *TEAD3* - involved in organ size control (and therefore may have a crucial role in determining the size of heart and lung for maximising oxygen utility at high altitude); *TULP1* – has photoreceptor function (possibly important for adaptation to intense light and UV radiation stress at high altitude); and *FKBP5* – an important modulator of stress response (including acute stress) due to its crucial role in the regulation of the hypothalamic-pituitary-adrenal (HPA) axis. Similarly, in E2, with minTemp contribution of 76%, the strongest *iHS*-sweep overlaps the *DDIT3* gene which is a multifunctional transcription factor in endoplasmic reticulum and plays an essential role in the response to a wide variety of cell stresses.

We also find genes *AGO4* and *CLSPN* from E9 and *SCARNA7* and *TRIM59* from E10 *iHS*- SRs; both ecotypes with a major environmental contribution from precSeasonality (84% and 80% respectively). These ecotypes show major variation in rainfall between dry (∼10 mm/m^2^) and wet seasons (>400 mm/m^2^). *AGO4* is involved in the gene silencing pathway and has been found upregulated during drought conditions in plants, *CLSNP* is involved in DNA repair, *SCARNA7* has role in RNA methylation (and thereby has modulatory effect on expression of genes) and *TRIM59* is involved in innate immunity. In ecotypes where multiple environmental variables have large contributions, we find genes involved predominantly in nervous system development/processes and transcriptional regulation, possibly to facilitate a concerted regulation of a wide variety of physiological functions.

While the above examples show genes from *iHS* analysis, the strongest *Hp*-based SRs were often detected from many ecotypes, possibly representing old sweeps. For instance, the strongest signals from E1-E3, E5-E7 and E10 overlapped with either the *TSHR* gene or its linked gene, *GTF2A1*. For some other ecotypes, we find genes involved in neurological processes (e.g., *EFHC2* in E8, *BEGAIN* in E9) and immune system development (*CACNA2D3* in E12). The strongest *Hp*-signal from E4 overlaps with several genes: *GRK7* – with a role in visual perception, *RNF7* – involved in protein ubiquitination pathway and response to redox state, and *ATPIB3* which serves as an ion pump across the plasma membrane that is essential for transepithelial transport including nutrient uptake. These genes appear highly relevant for adaptation to the major environmental drivers of the E4 ecotype – LandUse (49%) and SoilOrgC (43%), with both serving as proxies of food availability for scavenging chickens which require the ability to find food and assimilate available nutrients (Gheyas *et al*., 2021).

### Redundancy analysis (RDA) associates candidate sweeps with agro-climatic variables

As a multivariate linear regression approach, RDA allows simultaneous interrogation of many response variables (SNP genotypes) with many predictors (environmental variables) (Capblancq *et al.,* 2018). RDA was performed using 44,716 LD-pruned SNPs (∼9% of the total number of SNPs) overlapping the SRs.

The RDA model showed highly significant (*P* value = 0.001) deviation from the null hypothesis (no linear relationship between the SNP data and the environmental predictor). The model explained 1.9% of the genetic variance indicating that only a small proportion of the SR-SNPs have an association with environmental parameters. Each of the six RDA axes explained 10% to 30% of the variance captured by the model (Figure 7A). Only the first five axes were significant (*P*-value ≤ 0.01), and these together accounted for 90% of the total captured variance. The SNP loadings (Figure 7B) from these 5 axes were used to determine outliers i.e. SNPs showing significant association with environment.

**Figure 7.**
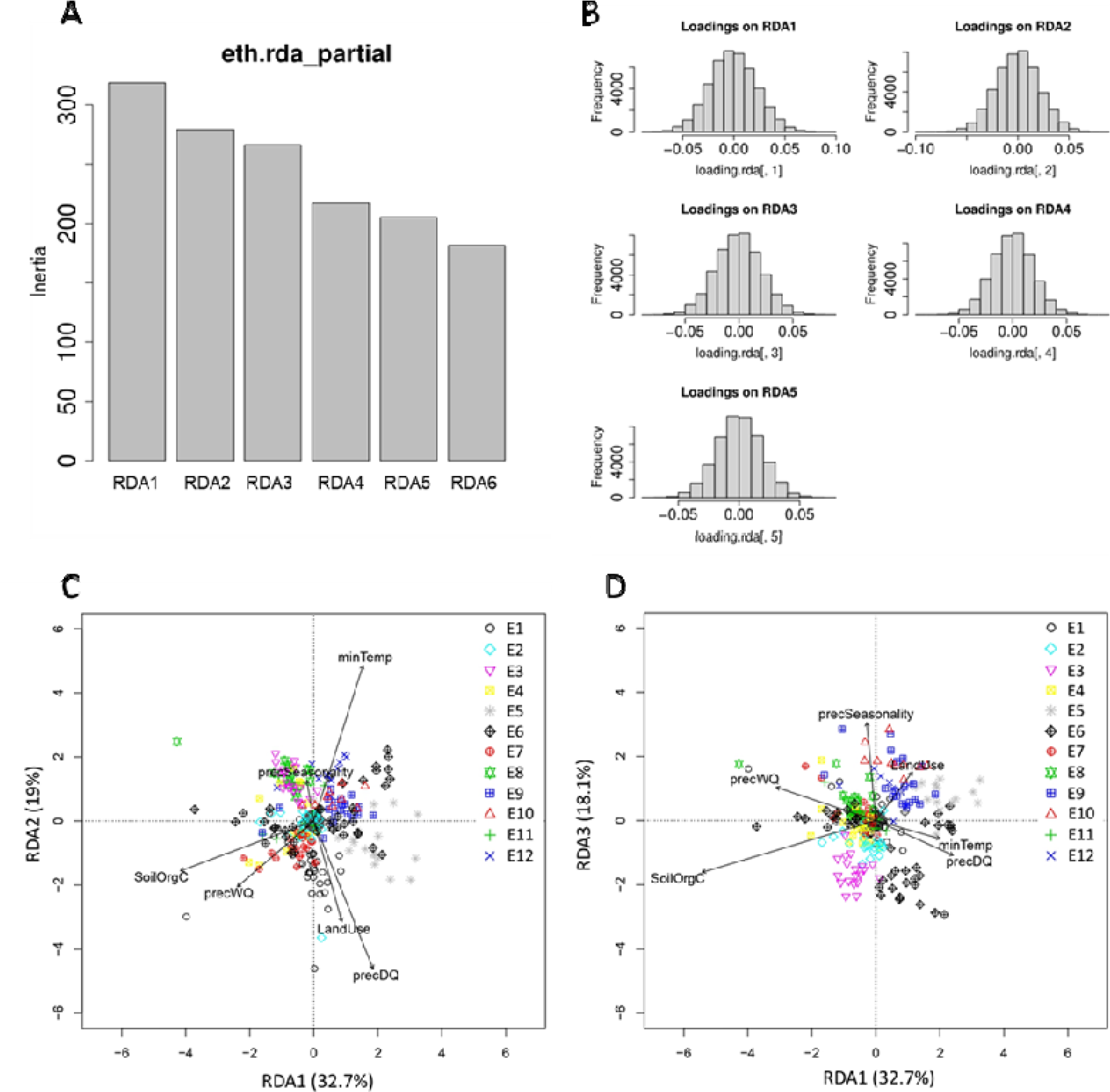
Results from RDA analysis (A) Projection of SNPs and environmental variables into the RDA space RDA1-RDA2 and (B) RDA1- RDA3; (C) Variance (inertia) explained by the six RDA axes; (D) Distribution of SNP loadings for significantly constrained axes (RDA1 – RDA5).

RDA plots show the distribution of the chicken samples from different ecotypes in relation to the RDA axes which are a linear combination of the predictor variables (Figures 7C-D). Some interesting relationships can be identified. For example, Figure 7C (axis1 vs axis2) shows that E1 genotypes are positively related to precDQ and LandUse, whereas E3 and E8 are negatively correlated with these two variables. Similarly, E12 genotypes are positively correlated with minTemp and negatively correlated with precWQ, whereas an opposite scenario for E7 is observed. By contrast, E9 individuals are positively associated with LandUse in Figure 7D (axis1 vs axis3).

Interestingly, individuals from E6 are dispersed in subgroups in both RDA plots. E6 comprises four different populations (Mihquan, HadushAdi, Hugub, and Jarso), and this is indeed one of the few ecotypes where the clustering based on niche overlap and Pearson correlation gave ambiguous results. These populations were included in the same ecotypes as all pairs showed values ≥ 0.6 for both the similarity metrics . For other ecotypes, no sub- clustering is evident when RDA plots are explored, confirming the overall robustness of our ecotype delineation. Regarding the E6, however, we need to keep in mind the limitation of RDA which can capture only linear association between genotype and environment. Therefore, in showing the distribution of the samples in the ordination space, RDA did not consider any non-linear association due to GxE interaction. ENM, on the other hand, is blind to any specific genotype-environment association; instead its characterisation is entirely dependent on the environmental conditions of the populations with the assumption that similar environmental pressure will lead to similar selective pressure on the genome. ENM-based ecotype characterisation is therefore expected to capture selective pressure from all sources (linear and non-linear).

Given a normal distribution of the SNP loadings in all axes (Figure 7B), SNPs that exceeded SD > 3 (*P*-value = 0.0027) in both tails were extracted as outliers. With this threshold, 616 SNPs (1.4% of total 44,716 SNPs tested) were found as outliers (Supplementary Table S9). The number of outliers varied in relation to the strongest correlated environmental variable, from 72 for precSeasonality to 130 for SoilOrgC. The strength of the environmental correlation of the outlier SNPs was generally low to moderate (*r* ≈ 0.1 – 0.4) (Supplementary Figure S8). The maximum correlation value was 0.42, identified for precWQ.

To gain an understanding of the biological functions of the genes associated with the outlier SNPs in relation to the correlated variables, a closer investigation was made on 30 RDA- outliers - which showed relatively large environmental correlation coefficients (*r* ≥ 0.3 for most predictors and ≥ 0.29 and ≥ 0.28 for PrecDQ and LandUse respectively as no outliers for these passed the first threshold) (Table 2). Notably, all these SNPs came from *iHS*- detected SRs. Highly relevant gene function or phenotypic associations are observed in most cases. For example, *PANK2* gene was detected in association with minTemp (or elevation). This gene is involved in neuronal and vascular development and respiration – important functions for thermo-tolerance and stresses at high-altitude. Two genes, *LDLRAD3* and *GPX7,* detected in association with precipitation variables (precSeasonality and precDQ), were previously reported in association with heat stress in chicken. These results are in agreement as the ability to cope with heat stress relies not only on the water availability but also on the environmental humidity – both factors being affected by rainfall patterns. *GPX7* has also been found associated with drought resistance in plants (Cruz de Carvalho, 2008). *LDLRAD3,* in our study, has also been detected in association with SoilOrgC - a proxy of scavenging conditions and food availability for chickens. Interestingly this gene has been found associated with lifespan expansion in a previous mouse study (Tyshkovskiy *et al.,* 2019).

In association with precipitation variables we also find several genes which are candidates for disease response, e.g. *MAPK4* – a candidate for *Salmonella* resistance in chicken, *LDLRAD3 –* associated with Newcastle Disease Virus infection in chicken, and *TNIP2 –* involved in immunity and inflammatory responses (Table 2). Since the rainfall pattern affects the prevalence of various pathogens, these detections appear very relevant. Furthermore, some genes detected in association with climatic factors (i.e., temperature and rainfall variables) are involved in broad regulatory and stress response pathways such as *MARCHF6* and *DET1* - protein ubiquitination and degradation pathways, *LRP6* – Wnt Signalling Pathway, *ERGIC3* – Endoplasmic Reticulum stress induced cell death and cell growth process, *CTBP1* – co-repression of diverse transcription regulators, and *RDH10* – Retinoid (Vitamin A) metabolic process with implications in many physiological processes including growth, development, immune system, and reproduction. On the contrary, the genes detected in association with SoilOrgC and LandUse (both potentially affect the nature and availability of food for foraging chickens as well as their foraging ability) show involvement in fat and carbohydrate metabolic processes (*LMF1, ADCK1, SLC2A6, OSBPL3*), growth, development and reproductive success (*GPCPD1, NHLRC2, THSD7B*), and neurological development (*PTPRZ1, AUTS2, NPAS3*).

The 616 RDA outliers intersected 349 SRs (33%), thereby providing indication of the major environmental variables exerting selection pressure on these specific SRs. The overlap of only a minority of SRs with the RDA-outliers can be explained by the fact that RDA identified only linear genotype-environment association across all samples. Any sweep regions resulting from a complex GxE interaction - thereby showing non-linear association - will not be detected by RDA. Outlier SNPs associated with all six environmental variables overlapped SRs from most ecotypes, confirming the combined contribution of different variables in each ecotype and thereby justifying the use of ENM for ecotype delineation and adaptation analysis. Also, some of the SRs overlap with several outliers associated with different agro-ecological variables. For instance, the SR on chr1:132930000_132960000 overlaps with RDA-outliers associated with SoilOrgC, minTemp and precDQ. This indicates that we are detecting the interaction of environmental variables that are collectively driving selection pressure on specific genomic regions. Such observation again highlights the importance and utility of using ENM-defined ecotypes for adaptation analysis.

### Regressing ecotype allele frequency of RDA-outliers with environmental predictors shows non-linear trends

The RDA analysis was performed to directly correlate SNP genotypes from individual samples with environmental parameters, without considering the ecotype effect. We wanted to further investigate how the ecotype allele frequencies of the outlier SNPs fluctuate with the ecotype average of the correlated environmental variables. The aim of this investigation was to gain an insight into whether the allele frequency shows a linear or non-linear trend with environmental variation. The investigation was made on the 30 strongest RDA-outliers (5 per environmental variable) by fitting linear and non-linear trend lines in scatter plots of allele frequency against ecotype average of environmental variables (Supplementary Table S10). Figure 8 shows scatter plots of 5 example SNPs (one for each variable). These plots as well as Supplementary Table S10 show that, in all cases, non-linear regression generated a larger R2 value i.e. better fit to the data. This result corroborates our assumption that ENM can capture complex GxE interaction in delineating ecotypes.

**Figure 8.**
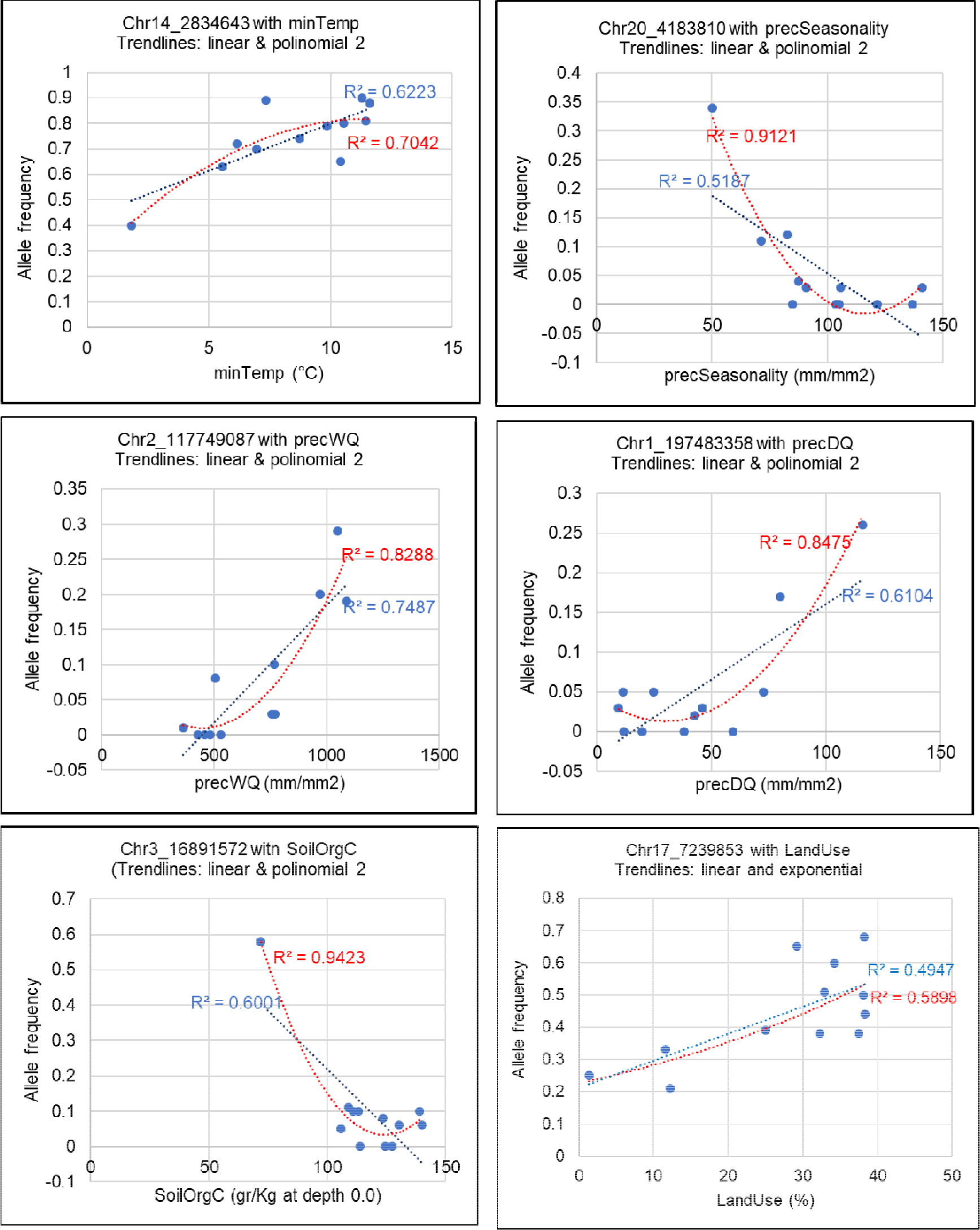
Allele frequency fluctuation of example RDA outliers with agro- climatic predictors (mean value) across ecotypes.One outlier (with the strongest RDA *r*-value) per predictor is shown as an example.

## Discussion

By integrating interdisciplinary approaches – ecological modelling with genomics - this study presents a novel framework for the identification and characterisation of indigneous livestock ecotypes showing genetic adaptation to distict agro-climatic conditions. Exemplified here with Ethiopian village chickens, the framework is fully transferable to any other livestock species, with important implications for conservation of adaptive biodiversity and breeding improvement towards achieving climate resilience. Unlike traditional adaptation studies, our approach directly models landscape heterogeneity based on a large specturm of environmental variables, thereby providing the opportunity to identify the key environmental pressures in livestock ecologies as well as capture the complex interplay of major variables in driving adaptive evolution at the genome level.

The benefits of our approach are visible at different levels. First, our method offers an opportunity to classify otherwise nondescript indigenous livestock populations into potential ecotypes based on a detailed characterisation of their agro-ecologies. As demonstrated in our results, classical agro-ecological zones are often insufficient in classifying livestock populations into ecotypes as those either lack resolution and precision or do not reflect the most relevant environmental predictors for the species in question. Contrarily, our method, based on ENM, offers not only the resolution and species specificity but also flexibility to incorporate any number of environmental variables in the characterisation process.

Another major advantage of our approach is its ability to capture complex interactions of many environmental variables, including the GxE interaction in shaping the livestock genome. This is possible through the use of different feature classes during the execution of ENM; e.g., we used 3 FCs: Quadratic (variance), Product (covariance i.e. captures interactions) and Hinge (linear response) (Phillips and Dudı’k 2007; Merow *et al*., 2013). The consequence of this is reflected in selective sweep detection. We find that even though the same variables may have a major contribution in defining multiple ecotypes (e.g., E9 and E10 have precSeasonality as the major driver with >80% contribution), the same candidate sweeps were not detected (only 10% of the candidate sweep windows are shared between these ecotypes), indicating possible interaction of other drivers in shaping adaptive evolution. Predicting GxE interaction is challenging and failure to take this into consideration has resulted in poorer performance of improved breeds in environmental conditions different from their original performance setting (Wakchaure *et al*. 2017). ENM-based characterisation of the agro-ecologies offers the interesting opportunity to assess the most suitable condition for any breed or population across a landscape. This has recently been tried for predicting suitable agro-ecologies for the introduction of exotic chicken breeds in Ethiopia (Lozano-Jaramillo *et al.,* 2018).

While most of the currently available methods of environmental association analysis (EAA) in landscape genomics – like RDA – capture only linear correlation between genotype and environment (Rellstab *et al*., 2015), our approach of integrating ENM with selective sweep analysis allowed capture of both linear and non-linear genetic responses to environmental pressure. That non-linear association plays a major contribution in driving adaptive evolution is reflected by the fact that we only detected a low to moderate level of correlation in the RDA analysis and found only one-third of the detected sweep regions to overlap with RDA outliers. Moreover, a basic inspection of the fluctuation of allele frequency of RDA outliers with environmental variables (Figure 8, Supplementary Table S10) demonstrated a greater power of non-linear regression in explaining variance compared to the linear approach. Modelling non-linear response is not yet well developed (Rellstab *et al*., 2015). The few available methods allowing non-linear EAA, include SAM (Joost *et al*., 2007) and

SAMβADA, which apply logistic regression methods. These methods only allow testing association of the presence/absence of an allele with environmental variables where interpretation of heterozygous genotypes becomes difficult. Moreover, unlike RDA, these are univariate analyses, allowing testing of only one genetic marker at a time, which fails to account for covariation among environmental variables and/or genetic markers (Capblancq and Forester, 2021). Therefore, no attempt was made to apply non-linear EAA in the present study.

In respect of genomic analysis, our approach has some added benefits of reducing false discovery rate (FDR) from the confounding effect of demography (e.g., genetic drift, founder effect, etc.). This is because multiple populations have been clustered together into most ecotypes, thereby increasing heterogeneity across the genome, except around the loci under selection pressure. To minimize FDR, we also employed a stringent criteria for identifying sweep signals. Instead of taking only the top 1% windows (empirical *P* value <0.01), as is frequently applied in selection siganture analyses, we employed further filtration based on standardised score (or Z score) with the same threshold applied to all ecotypes. This has resulted in a large variations in detected candidate sweeps from different ecotypes, indicating differential selection pressures.

One of the limitations of our study is that we only used chicken populations that were available to us. Although the sampling was performed to represent major Ethiopian AEZs, no specific environmental-gradation approach was applied to inform the sampling, as has been done in a recent paper by Kebede *et al*., (2021). Consequently, though fully valid, our study may not have surveyed all possible agro-climatic clines and the ecotypes presented here may not be an exhaustive list from Ethiopia. Although our study has considered a large array of environmental data, it could not incorporate some other potentially important drivers of adaptation, e.g., pathogenic or parasitic data, due to lack of publicly available data on such prevalence. Our approach, however, offers the exciting opportunity for such inclusion in future analyses. Indeed by integrating ecological concepts with genomics, our method opens up opportunities for interdisciplinary research. This approach can be employed to study the impact of climate change on indigenous livestock populations or to predict disease predisposition based on environmental conditions. This methodology will therefore have important implications for sustainable farming through climate resilience and conservation programs oriented to small-holder farmers, relying on local ecosystem production.

## Acknowledgments

This research was funded in part by the Bill & Melinda Gates Foundation (BMGF) and with UK aid from the UK Foreign, Commonwealth and Development Office (Grant Agreement OPP1127286) and was carried out under the auspices of the Centre for Tropical Livestock Genetics and Health (CTLGH), established jointly by the University of Edinburgh, SRUC (Scotland’s Rural College), and the International Livestock Research Institute. The findings and conclusions contained within are those of the authors and do not necessarily reflect positions or policies of the BMGF nor the UK Government. This research was conducted as part of the Consultative Group on International Agricultural Research (CGIAR) Research Program on Livestock and is supported by contributors to the CGIAR Trust Fund. The study constituted part of Adriana Vallejo-Trujillo’s PhD research that was funded by Vice-Chancellor Scholarship for Research Excellence International at University of Nottingham and Administrative Department of Science, Technology and Innovation (Colciencias) – Colombian Government (Call 2015 N°728). We thank Prof. Nick Sparks (CTLGH, SRUC) for his valuable support in conducting this research. We would also like to thank Edinburgh Genomics (Edinburgh, UK) for producing the sequence data used in this study.

## Data Accessibility

The whole genome sequence data used in this study is publicly available via the project accession number PRJEB39275 in European Nucleotide Archive (ENA) and the SNP data is available via European Variation Archive (EVA) with the accession numbers, Project:PRJEB46494

## Author contributions

OH, AG, AVT and JS conceived the research project. AK, TD, and OH led the collection of samples and population metadata. MLJ contributed with the ENM approach delineation. AVT and AG performed the analyses and led the writing of the manuscript but all authors contributed critically to the drafts.

## Supplementary Tables

Supplementary Table S1. Twenty five Ethiopian village chicken populations studied and their agro-climatic conditions

Supplementary Table S2. AUC values from individual population models - for overall model and for individual environmental variables

Supplementary Table S3a. Heat map of pairwise Niche Overlap (I) statistics between suitable maps (logistic scale) of 25 Ethiopian chicken populations

Supplementary Table S3b. Heat map of pairwise Pearson correlation (r) statistics between suitable maps (logistic scale) of 25 Ethiopian chicken populations

Supplementary Table S4: Candidate selection signature regions (SRs) and overlapping genes detected from analyses of different ecotypes

Supplementary Table S5. Candidate sweep regions commonly detected by Hp and iHS either from the same ecotype or from different ecotypes.

Supplementary Table S6a. Genes from Hp-based candidate sweeps regions from different ecotypes

Supplementary Table S6b. Genes from iHS-based candidate sweeps regions from different ecotypes

Supplementary Table S7: Sweep regions overlapping chicken QTLs

***Supplementary Table S8a: Genes overlapping the strongest iHS signals from each ecotype and their functional relevance to the environmental condition of the corresponding ecotype*.**

Supplementary Table S8b: Genes overlapping the strongest Hp signals from each ecotype and their functional relevance to the environmental condition of the corresponding ecotype.

Supplementary Table S9: Identified outlier SNPs by applying RDA on Hp and iHS SNPs of candidate regions (SD > 3 and P-value = 0.0027 two-tailed cut-off)

***Supplementary*** Table S10. ***Fluctuation of ecotype allele frequency with mean environmental value of 30 strongest RDA outlier SNPs (5 per environmental variables)***

Supplementary Figures

Supplementary Figure S1: The relative contribution of the 6 selected environmental variables (in terms of permutation importance) in individual chicken populations.

Supplementary Figure S2: Suitability maps (in logistic output) of individual chicken populations across the Ethiopian landscape based on 6 environmental variables.

Supplementary Figure S3: The relative contribution of the 6 selected environmental variables (in terms of permutation importance) in individual ecotypes.

Supplementary Figure S4a: Correlation between Hp and mean unstandardized iHS values by ecotype

Supplementary Figure S4b: Correlation between Z(Hp) and mean standardized iHS values by ecotype

Supplementary Figure S5. Proportion of detected sweep regions by iHS, Hp, and iHS-Hp in different ecotypes

Supplementary Figure S6. Percentage of shared selective sweep windows (SSWs) among ecotypes

Supplementary Figure S7: Functional classification of genes overlapping sweep regions in different ecotypes according to Panther biological pathways. The numbers shown on the right hand side indicate the number of ecotypes where the pathway is represented.

Supplementary Figure S8. Boxplots summarizing the environmental correlation of RDA outlier SNPs. Y axis shows the absolute values of correlation coefficient

