## Supplementary Figure for "A framework for defining livestock ecotypes based on ecological modelling and exploring genomic environmental adaptation: the example of Ethiopian village chicken"

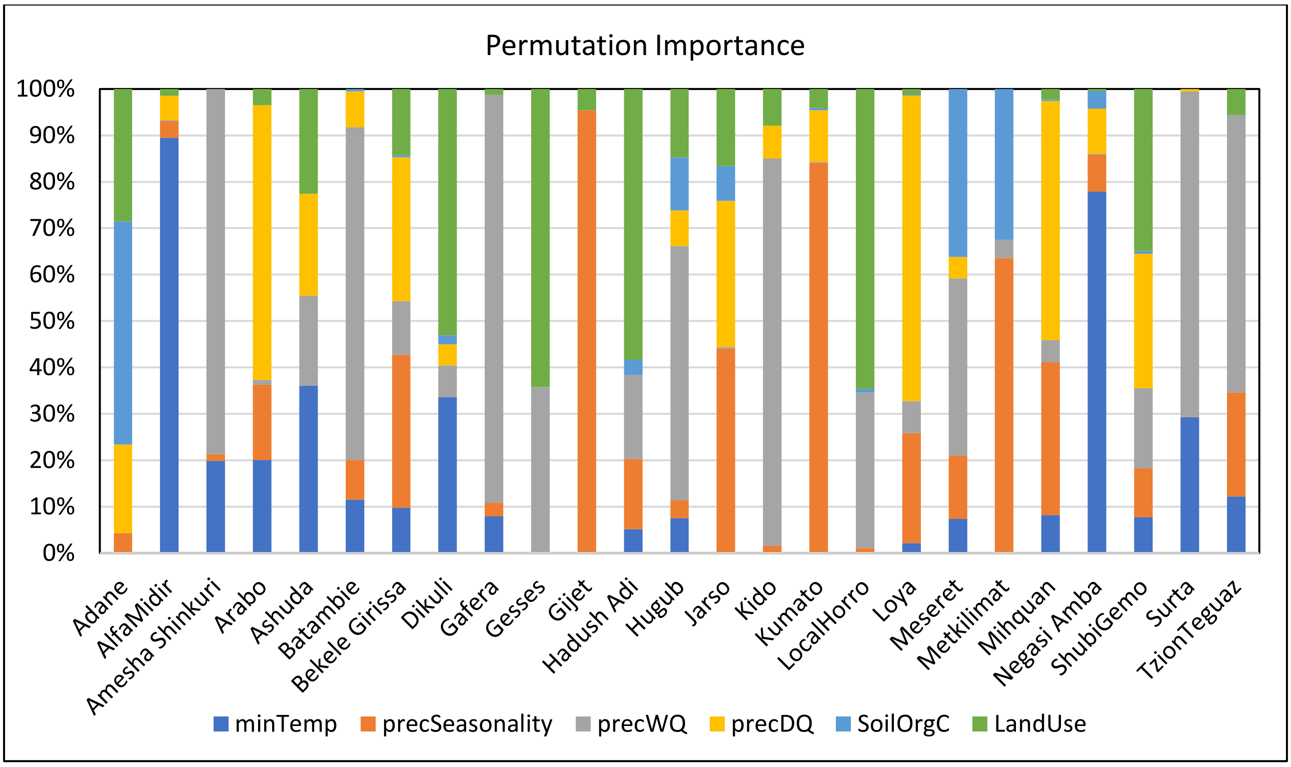


**Supplementary Figure S1: The relative contribution of the 6 selected environmental variables (in terms of permutation importance) in individual chicken populations.**

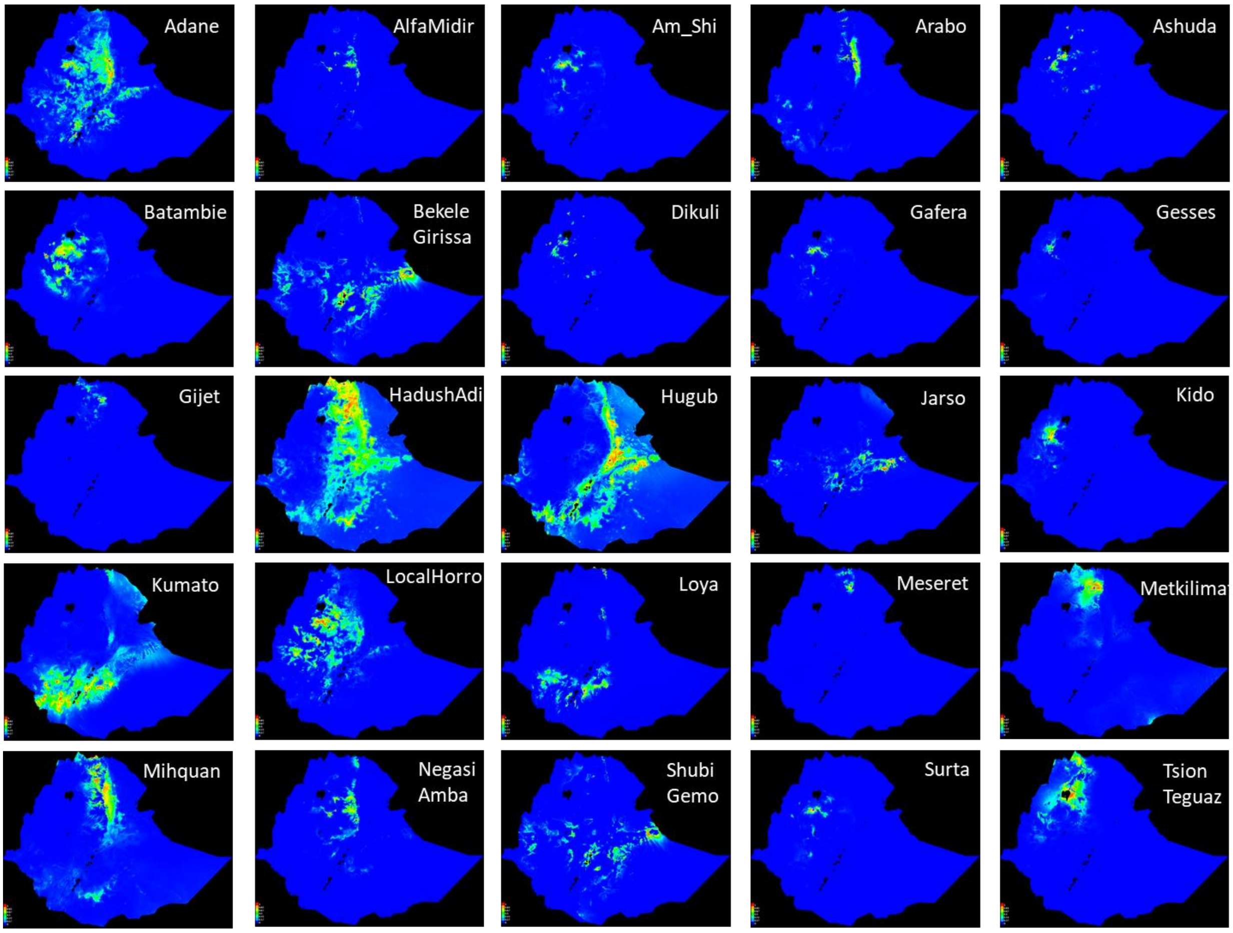


**Supplementary Figure S2: Suitability maps (in logistic output) of individual chicken populations across the Ethiopian landscape based on 6 environmental variables.**

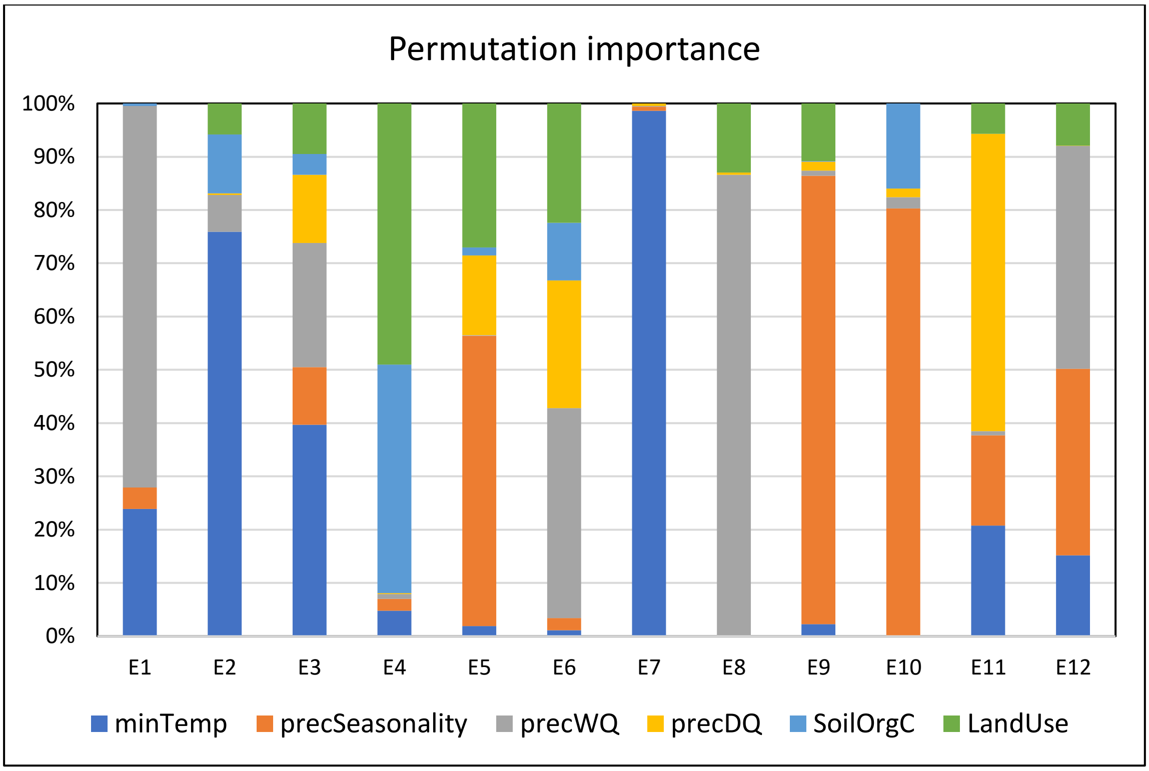


**Supplementary Figure S3: The relative contribution of the 6 selected environmental variables (in terms of permutation importance) in individual ecotypes.**

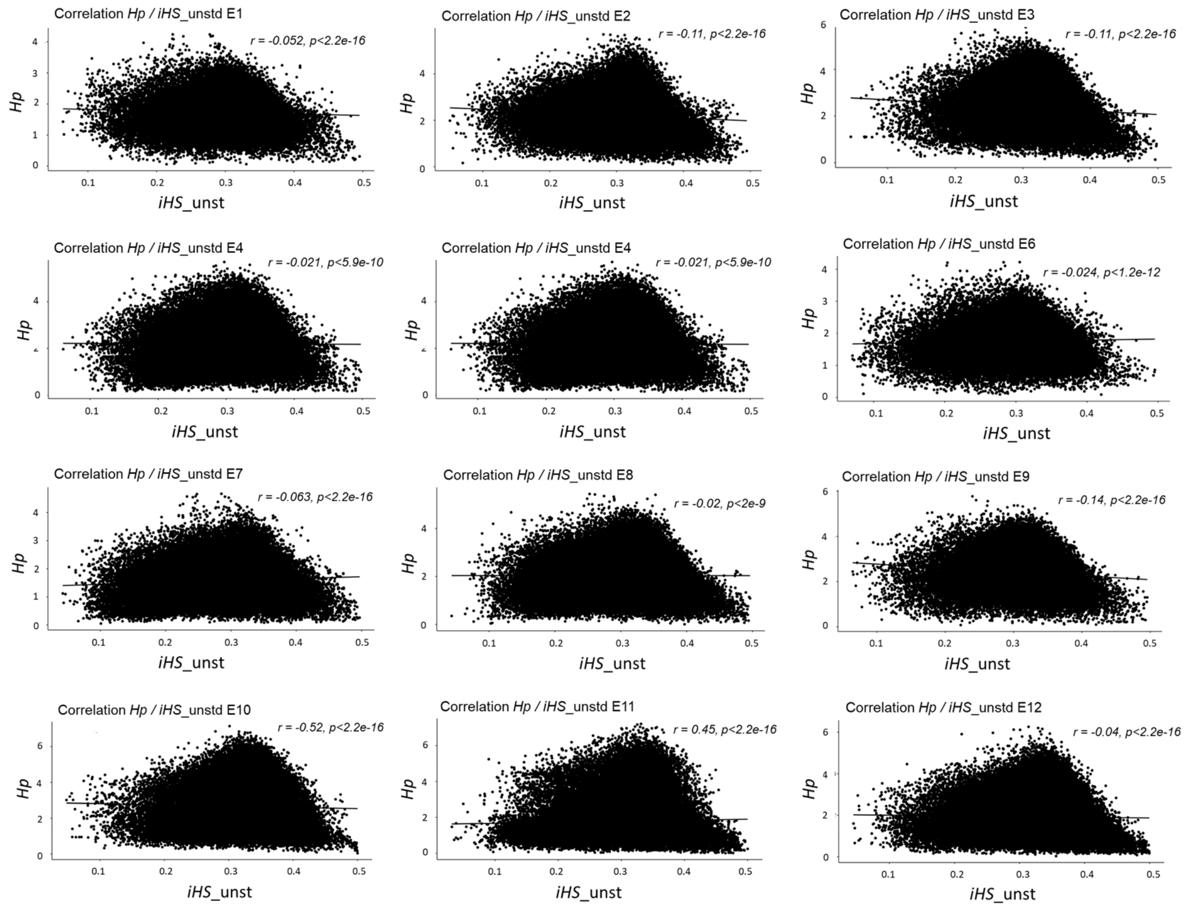


**Supplementary Figure S4a: Correlation between *Hp* and mean unstandardized *iHS* values by ecotype**

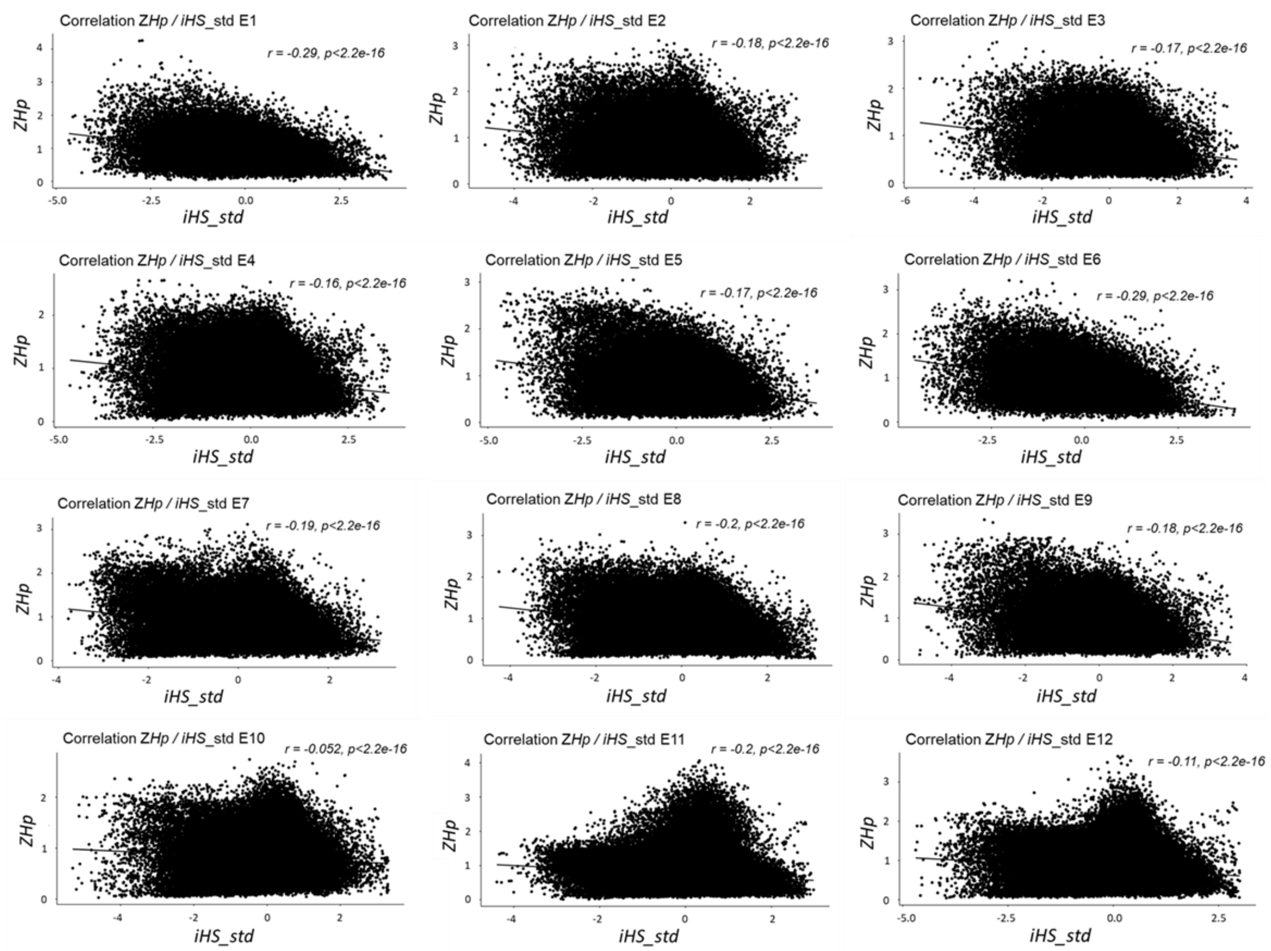


**Supplementary Figure S4b: Correlation between *Z(Hp)* and mean standardized *iHS* values by ecotype**

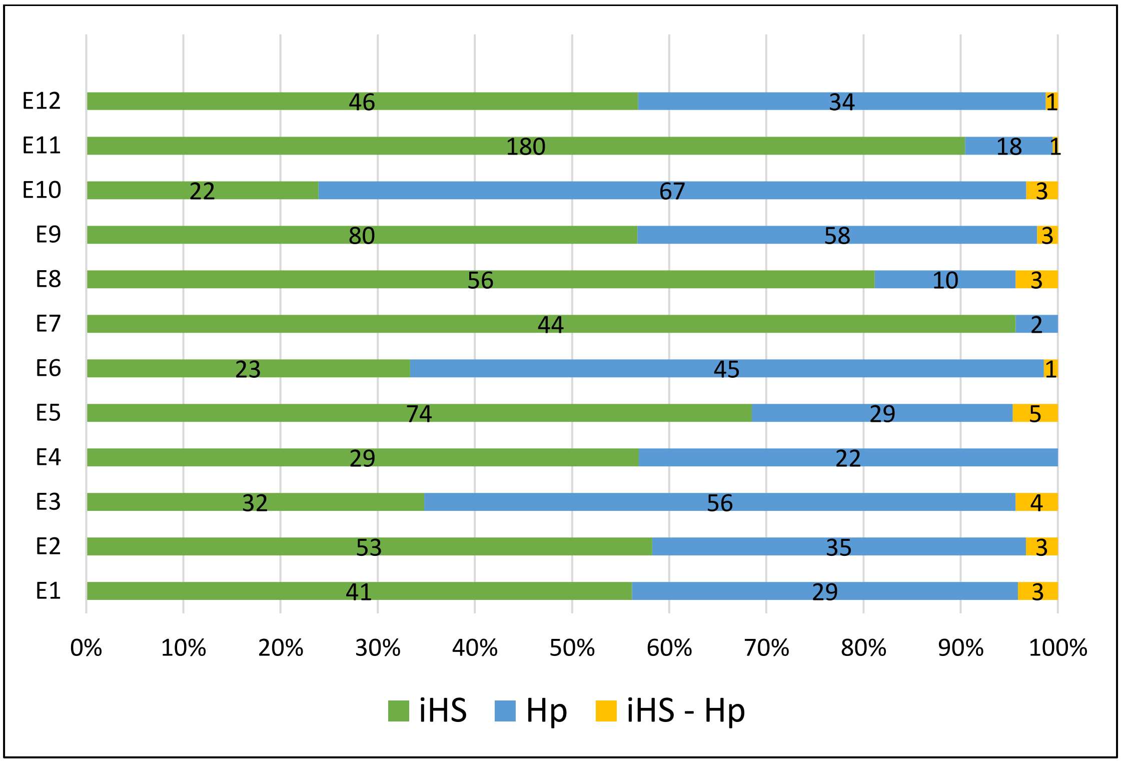


**Supplementary Figure S5. Proportion of detected sweep regions by *iHS*, *Hp*, and *iHS-Hp* in different ecotypes . (‘*iHS – Hp’* indicates sweeps that were simultaneously detected by both methods and some of them are common across ecotypes).**

| #SSWs | 300 | 354 | 368 | 225 | 549 | 224 | 212 | 327 | 689 | 355 | 608 | 300 |
|---|---|---|---|---|---|---|---|---|---|---|---|---|
| | E1 | E2 | E3 | E4 | E5 | E6 | E7 | E8 | E9 | E10 | E11 | E12 |
| E1 | | | | | | | | | | | | |
| E2 | 7 | | | | | | | | | | | |
| E3 | 7 | 6 | | | | | | | | | | |
| E4 | 5 | 6 | 10 | | | | | | | | | |
| E5 | 5 | 5 | 11 | 9 | | | | | | | | |
| E6 | 9 | 4 | 10 | 9 | 8 | | | | | | | |
| E7 | 2 | 2 | 2 | 6 | 2 | 4 | | | | | | |
| E8 | 4 | 6 | 3 | 6 | 6 | 2 | 2 | | | | | |
| E9 | 5 | 4 | 12 | 3 | 8 | 6 | 2 | 3 | | | | |
| E10 | 7 | 6 | 13 | 5 | 10 | 8 | 1 | 5 | 10 | | | |
| E11 | 1 | 2 | 1 | 1 | 2 | 1 | 1 | 1 | 1 | 1 | | |
| E12 | 7 | 4 | 6 | 6 | 7 | 7 | 2 | 4 | 5 | 8 | 1 | |

**Supplementary Figure S6. Percentage of shared selective sweep windows (SSWs) among ecotypes**

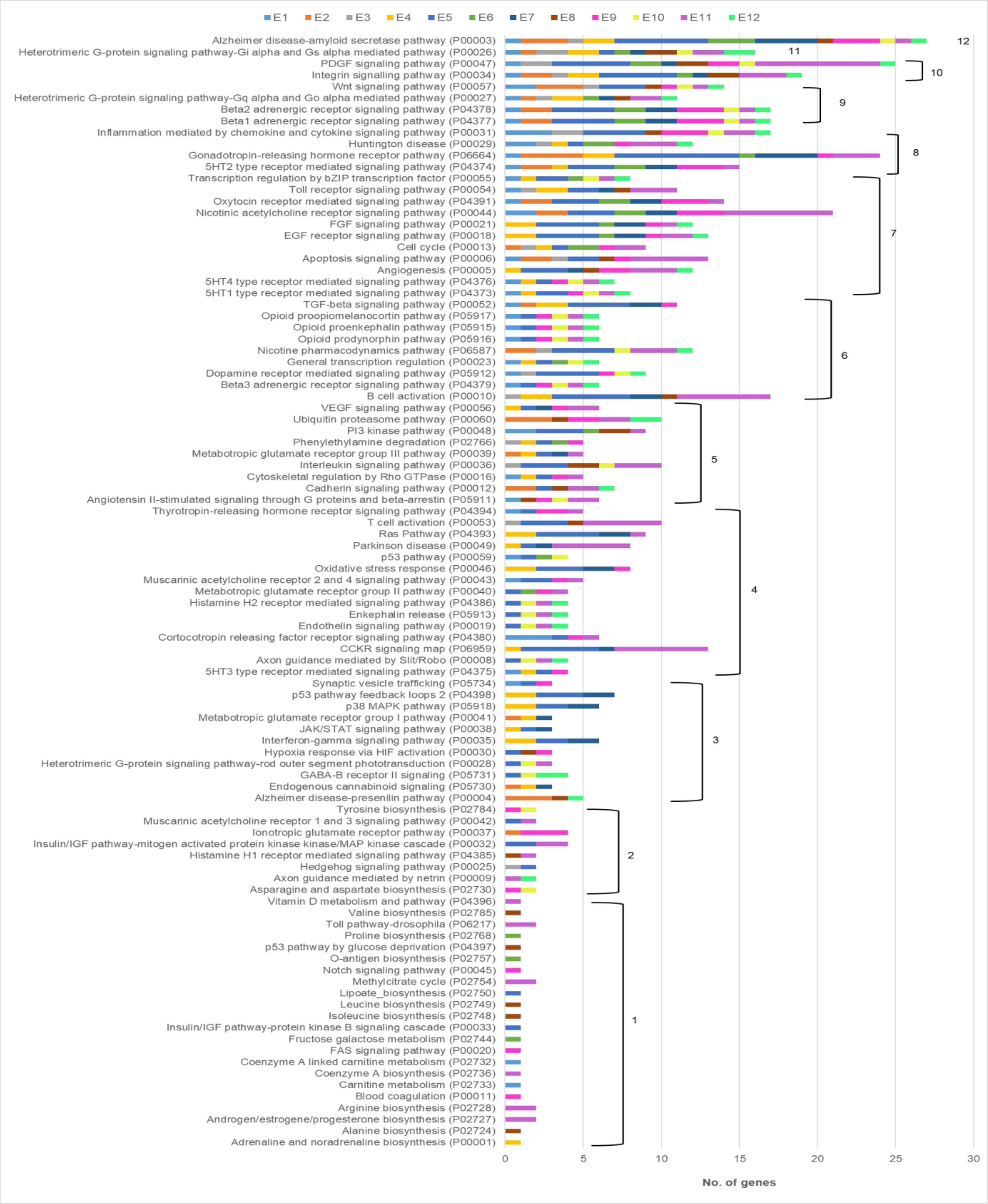


**Supplementary Figure S7: Functional classification of genes overlapping sweep regions in different ecotypes according to Panther biological pathways. The numbers shown on the right hand side indicate the number of ecotypes where the pathway is represented.**

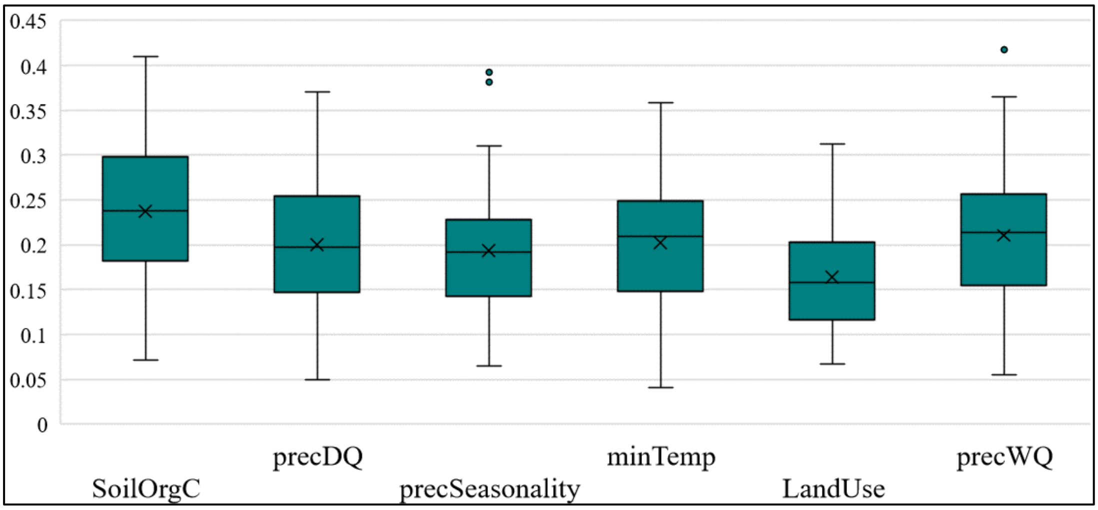


**Supplementary Figure S8. Boxplots summarizing the environmental correlation of RDA outlier SNPs. Y axis shows the absolute values of correlation coefficient**
